# Interlocus gene conversion causes mosaic divergence in tandem paralogues – modeling *HMA4* evolution in *Arabidopsis halleri*

**DOI:** 10.1101/2025.10.29.685298

**Authors:** Yannick Schäfer, Yichen Zheng, Piyal Karunarathne, Akash Chandra Parida, Björn Pietzenuk, Vinod Kumar, Laura E. Rose, Ute Krämer, Thomas Wiehe

## Abstract

Following gene duplication, the evolutionary trajectories of gene copies can be shaped by various processes including mutation, selection, recombination and interlocus, i.e., non-allelic, gene conversion (IGC). To explore dynamics and consequences of these processes, we developed a mathematical model that simulates the early evolution of recently duplicated, tandemly arrayed paralogous gene copies with positive selection on the new gene copy. We compared the results of our model to sequence variation in the three tandemly arrayed *Arabidopsis halleri HEAVY METAL ATPase 4* (*HMA4*) gene copies, which are known to undergo IGC. We find that the effective rates of IGC vary within genes, resulting in a mosaic pattern of divergence. Informed by empirical data, our model captures the impact of IGC on the diversity within each gene copy and the divergence among them. By tailoring the model to the *AhHMA4* gene copies, we demonstrate the model’s flexibility and its potential to provide insights into the evolutionary dynamics driving the evolution of tandemly arrayed paralogues. This study enhances our understanding of the balance between concerted evolution and divergence in the early evolution of gene families following gene duplication.

## Introduction

Genetic variation arises through a variety of processes. One of them is tandem gene duplication, which plays a key role in the generation of new genetic material (Cannon et al., 2004; English et al., 2023). This process can produce arrays of multiple paralogous copies of an original single gene and thereby increase the dosage of the gene product. Alternatively, and depending on the degree of divergence, paralogues may acquire partially or entirely different functions, undergo pseudogenization or may be lost again (Freeling, 2009; Hofberger et al., 2013). Tandem gene duplications occur in all kingdoms of life and can facilitate adaptation to changing or new environments (Qiao et al., 2019; Hanada et al., 2008).

Highly similar gene duplicates located in tandem in a genome are often subject to a process addressed as *interlocus*, *non-allelic* or *ectopic*, gene conversion (IGC). IGC occurs during the repair of DNA double strand breaks and entails the non-reciprocal transfer of a segment of several tens to hundreds of base pairs from one paralog to another (Klein and Petes, 1981; Gao and Innan, 2004; Mondragon-Palomino and Gaut, 2005). Signatures of repeated IGC among paralogues are high sequence similarity and an excess of shared polymorphisms (Stephens, 1985; Nagylaki, 1984; Dumont, 2015). Conversely, the frequency of IGC reduces with decreasing sequence similarity between donor and target sequences. A meta analysis of human pathogenic gene conversions showed that the homology between donor and acceptor gene sequences is always at least 92% and usually more than 95% (Chen et al., 2007). IGC is an important evolutionary factor in all kingdoms of life, from prokaryotes to fungi and plants (Morris and Drouin, 2004; Kupiec and Petes, 1988; Mondragon-Palomino and Gaut, 2005).

The HMA4 protein of *Arabidopsis thaliana* is a *P*_1B_-ATPase that mediates the export of Zn^2+^ and Cd^2+^ ions across the plasma membrane of cells in the vasculature (Hussain et al., 2004). It contributes to root-to-shoot translocation of these ions and its expression in shoots indicates that it could also mediate their distribution within the shoot. (Hussain et al., 2004; Verret et al., 2004). The *HMA4* gene cluster on chromosome 3 of *A. halleri* consists of three tandemly arrayed gene copies (Hanikenne et al., 2008). The *HMA4* locus in *A. halleri* has experienced two duplication events, previously estimated to have occurred around 357 kyr and 250 kyr ago (Roux et al., 2011). Based on sequence similarity, the recent duplication produced the genes *HMA4-2* and *HMA4-3* (Hanikenne et al., 2008). The three resulting gene copies are present in all accessions of *A. halleri* that were examined so far (Hanikenne et al., 2013). A reverse genetic approach demonstrated that the three tandem gene copies of *HMA4* are required for Zn and Cd hyperaccumulation and for the full extent of Zn and Cd hypertolerance, naturally selected extreme traits characteristic of the species *A. halleri* (Hanikenne et al., 2008). Metal hyperaccumulation is defined as the accumulation of extraordinarily high levels of metals of more than 10 times the tolerance limits of non-hyperaccumulating species in above-ground tissues in the natural habitat of a species, i.e. more than 3 mg Zn g^−1^ dry biomass and more than 100 µg Cd g^−1^ dry biomass (Krämer, 2010). More than 700 metal hyperaccumulator taxa have been identified to date. Found in plants native to soils containing particularly high levels of heavy metals of geogenic or anthropogenic origin, metal hypertolerance is an unusually high degree of tolerance to heavy metals far beyond the basal metal tolerance common to all organisms (Clemens, 2001; Krämer, 2010), and generally in metal hyperaccumulator plants (i.e., also when their habitat is a non-polluted soil). The combination of triplication and *cis-regulatory* mutations of *HMA4* in *A. halleri* results in a strongly elevated total transcript abundance of *HMA4* in both roots and shoots, compared to closely related species *A. thaliana* and *A. lyrata* (Hanikenne et al., 2008; Castanedo et al., 2025).

The *HMA4* locus in *A. halleri* has previously been described as a target of natural selection through a hard selective sweep affecting the promoter regions, coupled with an exchange of genetic material between coding sequences of the three copies via IGC (Hanikenne et al., 2013). As expected after a selective sweep, the *HMA4* locus showed drastically reduced sequence diversity and a negative Tajima’s *D* compared to flanking genomic regions, and this was the most pronounced in between and directly upstream of the coding sequences of the three *HMA4* gene copies, including their promoter sequences. However, this pattern of low sequence diversity was interrupted in the coding regions of the *HMA4* gene copies, which exhibited network-like sequence relationships, which the authors attributed to IGC. This earlier study was based on copy-specific amplicons designed according to an *A. halleri* BAC contig so that the analyzed short sequence segment was located at the 3’ ends of the *HMA4* coding sequences. In this study, we chose the *HMA4* genes of *A. halleri* as a model case for studying IGC between duplicated gene copies of recent origin taking advantage of sequence data from *de novo* genome assemblies.

Previous models employed to capture the relationship between IGC and divergence among gene copies were developed, for instance, by Teshima and Innan (2004) or Thornton (2007). Teshima and Innan (2004) used a forward-in-time approach focusing on the suppression of IGC once a divergence threshold is surpassed. However, their model did not include factors such as selection and demography. Thornton’s (2007) model is based on the coalescent process and takes into account population size and demography. He derived the site frequency spectra (SFS) of the original and the duplicated gene copies under variable recombination and IGC rates and showed that the newly duplicated copy exhibits a reduction in diversity compared to the ancestral gene copy and an excess of rare alleles. More recently, the forward-in-time simulation study by Hartasánchez et al. (2014) overcomes the limitation of considering only one pair of duplicated genes. They evaluated the effects of reciprocal recombination (crossing-over) and IGC on the distributions of single nucleotide polymorphisms and of linkage disequilibrium (LD). Recombination and IGC are shown to interact in a complex way; for example, IGC can break down LD patterns created by recombination. The existence of recombination hotspots can also lead to a heterogeneous distribution of variation across the duplicated locus (“patterning”). Still, because their simulation model assumed neutrality, neither the role of purifying nor that of positive selection was assessed.

Here, we implemented a forward-in-time simulation model of gene duplication in which IGC decreases with rising divergence. It considers (a) positive selection for the new copy (selection for enhanced dosage), (b) purifying selection on the coding sequences of all copies, and (c) a gradually diminishing rate of IGC as sequence divergence increases. We show that under such a model, coding regions – due to their functional constraints – may enter an IGC/mutation equilibrium while introns of the same locus continuously diverge. We thoroughly analyse the structure of the *AhHMA4* gene copies and the sequence variation at the respective loci. Subsequently, we complement these results with computer simulations based on our model to further explore the evolutionary forces at this locus and to re-examine its duplication history.

## Results

### Empirical Observations

#### Structure of the HMA4 genes in Arabidopsis halleri and its close relatives

The three tandemly arrayed *HMA4* gene copies of *A. halleri* are positioned in a head-to-tail arrangement (Hanada et al., 2008; Hanikenne et al., 2013) and contain 11 exons. Exon 0 and the initial nucleotides of exon I, as well as the 3’ part of exon IXb correspond to the 5’ and 3’ UTRs, respectively. The protein-coding part of the sequence spans from exon I to the 5’ part of exon IXb. The sizes of introns II and III (long introns) differ strikingly from those of introns IV to X (short introns). The protein is composed of a short cytosolic *N* -terminus encoded by exon I and the start of exon II, a transmembrane (TM) domain with eight TM helices encoded by exons II to IXa, and a long cytosolic *C*-terminus encoded by parts of the exons IXa and IXb (Figure 1).

**Figure 1:**
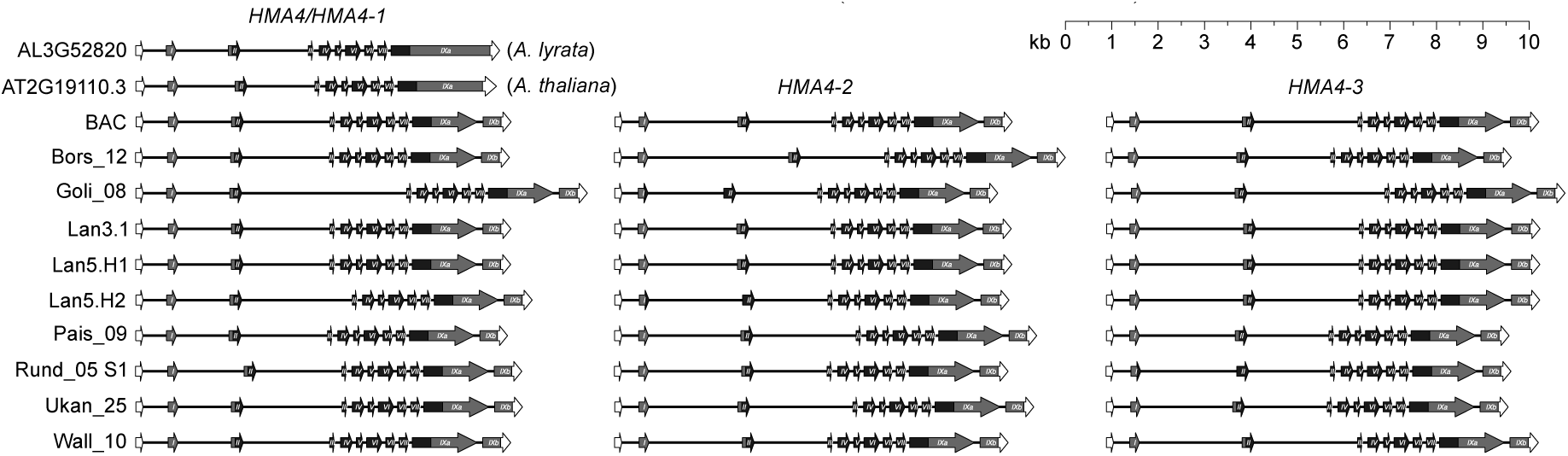
Intron/Exon structure of *HMA4* genes from different accessions of *A. halleri* compared to *A. lyrata* and *A. thaliana* (top left panel). Filled arrows mark exons in the 5’ to 3’ direction (left to right): **white:** 5’ untranslated region (UTR); **grey and black:** coding regions [grey: N-terminus; black: transmembrane (TM) domain-containing region; grey: cytosolic C-terminus]; **white:** 3’ UTR. The “BAC” (Bacterial Artificial Chromosome) sequences were taken from two BACs sequenced by Hanikenne et al. (2008) (Genbank accessions EU382072 and EU382073), for other designations of genotypes see Stein et al. (2017). All annotations are presented to scale, ensuring that the relative lengths of gene features are directly comparable across genes.

We extracted the *HMA4* locus from 9 high-quality *de novo* genome assemblies of different accessions of *A. halleri* (see Materials and Methods). While the exon-intron structure of all *HMA4* genes is conserved across all of these haplotypes, there are a few outliers in the lengths of the long introns. The most striking examples are intron III of *HMA4-1* & *HMA4-3* in individual Goli 08 and intron II of *HMA4-2* in individual Bors 12 (see Fig. 1)(Stein et al., 2017), both of which are much longer than the corresponding introns in the other accessions and *HMA4* gene copies.

When looking beyond species boundaries we find that the single copy genes AL3G52820 in *A. lyrata* and AT2G19110.3 in *A. thaliana* have a gene structure which is almost identical to that of the *A. halleri* copies. The only difference is an additional intron in *A. halleri*, which splits the last exon into two compared to the *A. thaliana* and *A. lyrata* homologs (Figure 1, Table S1).

#### Genetic variation in the HMA4 paralogues in A. halleri

*HMA4* sequences from *A. halleri* were sorted into the three categories ‘short introns’, ‘long introns’ and ‘CDS’ with short introns spanning up to 150 bp and long introns at least 1 kb (Table S1). The intermediate intron between the 5’ UTR and the first coding exon was not part of this analysis. Afterwards, the sequences were aligned by category and filtered to exclude non-aligning positions, via T-Coffee (Notredame et al., 2000). We then calculated the number of pairwise differences within and between gene copies for each type. For the CDS, we counted synonymous and nonsynonymous differences separately. We did not use a codon substitution model, but checked empirically for each pairwise comparison and each variable nucleotide pair whether the difference was synonymous or not. Counts were then scaled to numbers per bp. Boxplots of these numbers are shown in Figure 2. We also calculated for each gene and each type Watterson’s *θ* (Watterson, 1975), an estimator of the mutation rate scaled by effective population size. Discrepancies between *θ_w_* and the average of the pairwise differences (*θ_π_*) can be indicative of the presence of natural selection. As a baseline, Fischer et al. (2017) estimated a genome-wide average of *θ̅_w_ ≈* 0.0085 from 180 *A. halleri* individuals.

**Figure 2:**
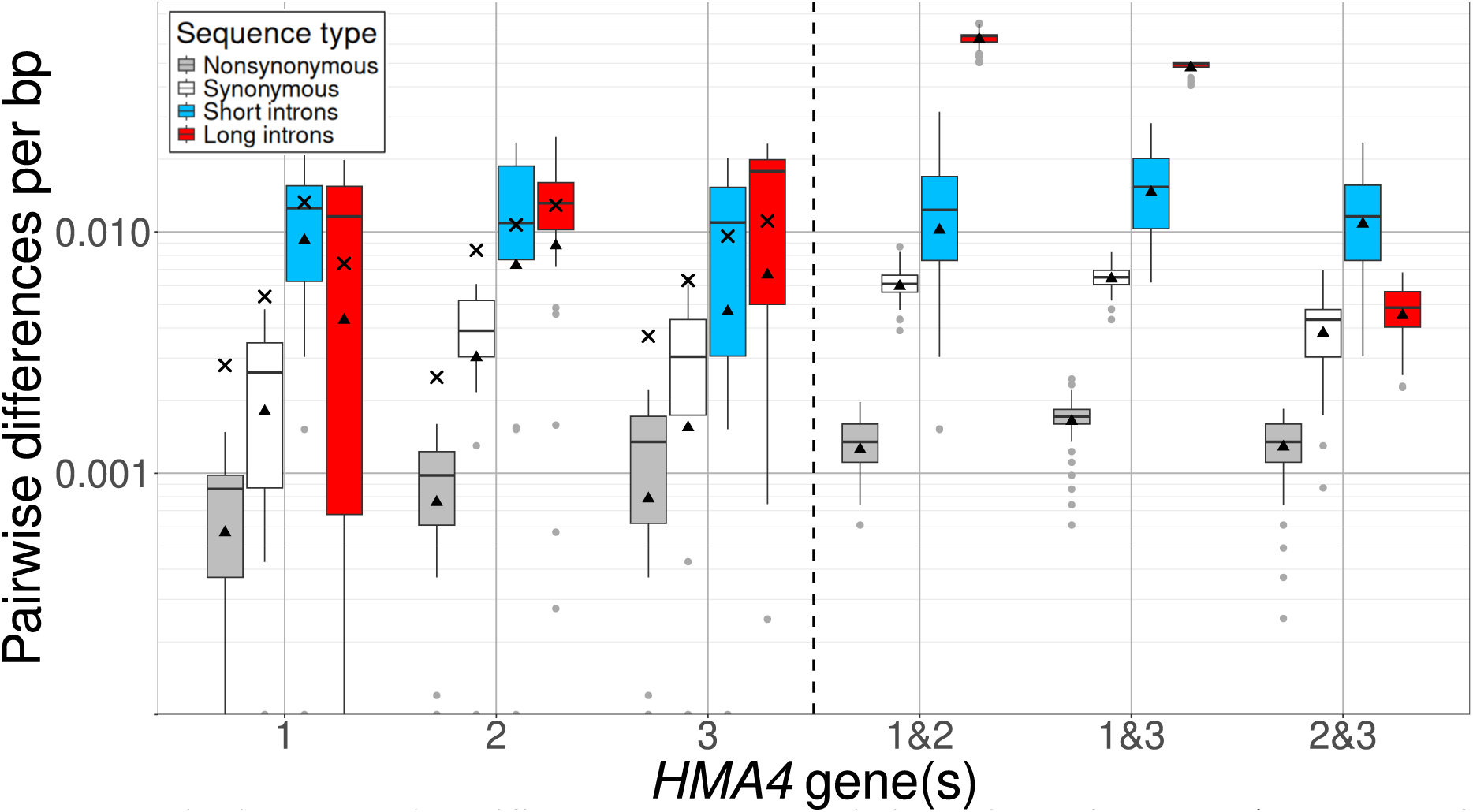
Pairwise nucleotide differences per base pair in regions of the *HMA4* genes. Left panel: average pairwise diversity (*θ_π_*) within individual gene copies. Right panel: average pairwise divergence (d) between gene copies (without correction for multiple hits). Calculations are based on multiple sequence alignments generated using `T-Coffee` and `mafft`. Introns II and III (see Figure 1 are classified as long introns (*>* 1 kb), while introns IV through X are classified as short introns (see Table S1 for details). The y-axis is log10-scaled.

We observe a nucleotide diversity (*θ_π_*) in all introns of roughly 1% in all three genes. At synonymous coding sites, *θ_π_* is around 0.3% and in nonsynonymous sites between 0.07% (in *HMA4-1*) and 0.12% (in *HMA4-3*) (Table 1), similar to diversity reported earlier in *AhHMA4* coding regions Hanikenne et al. (2013). The sequence divergence between gene copies is similar among all three between-gene comparisons when focusing on short introns (from 1.2% to 1.6%). In synonymous sites, we observe *θ_π_* between 0.4% (in 2&3) and 0.6% (in 1&2 and 1&3). In nonsynonymous sites, it is between 0.13% and 0.17%. The striking exception to this pattern of similarity of the values among the different gene pair comparisons are the long introns: the younger duplicates (*HMA4-2* and *HMA4-3*) have diverged much less (0.5%) from one another than either of them has diverged from *HMA4-1* (6.3% and 4.8% respectively), see Table 1. Moreover, while between-copy divergence exceeds within-copy diversity by a factor of up to 2 in short introns and coding exons, this factor is consistently around 5 in long introns. This observation fits the idea that the frequency of IGC decreases with increasing sequence divergence.

**Table 1:** Diversity and divergence in introns and CDS of *AhHMA4*.

| *HMA4*- | diversity within genes* |  |  |  |  |  | divergence between genes† |  |  |
| --- | --- | --- | --- | --- | --- | --- | --- | --- | --- |
|  | 1 |  | 2 |  | 3 |  | 1 & 2 | 1 & 3 | 2 & 3 |
| | $\theta_\pi$ | $\theta_w$ | $\theta_\pi$ | $\theta_w$ | $\theta_\pi$ | $\theta_w$ | | | |
| intr<sup>short</sup> | 0.0116 | 0.0133 | 0.0117 | 0.0107 | 0.0093 | 0.0096 | 0.0125 | 0.0156 | 0.0120 |
| intr<sup>long</sup> | 0.0103 | 0.0074 | 0.0120 | 0.0129 | 0.0128 | 0.0111 | 0.0632 | 0.0481 | 0.0047 |
| CDS<sup>syn</sup> | 0.0025 | 0.0054 | 0.0038 | 0.0084 | 0.0028 | 0.0063 | 0.0061 | 0.0064 | 0.0041 |
| CDS<sup>nonsyn</sup> | 0.0007 | 0.0028 | 0.0009 | 0.0025 | 0.0012 | 0.0037 | 0.0013 | 0.0017 | 0.0013 |

\* $\pi$ : average number of nucleotide differences per bp across $\sum_{n=1}^9 n = 45$ pairwise sequence comparisons
$\theta$ : number of polymorphic sites per bp within a gene divided by $h_{n-1} = 2.82897$ , where $n = 10$ is the sample size
† average number of nucleotide differences per bp across $\sum_{n=1}^{10} n = 55$ pairwise between-gene sequence comparisons

One effect of IGC is that polymorphisms which arise through mutation in one gene copy can be transferred to another gene copy. Without IGC, such shared polymorphisms occur very rarely because they require the independent mutation of the exact same position within two (or even three) gene sequences into the same alternative nucleotide. To visualize this effect among the *HMA4* gene copies, we compared the numbers of shared and private polymorphisms in the different gene regions (Figure 3, Table 2). In total, we observe 280 polymorphic sites in the three *HMA4* genes. Among these, 150 are private polymorphisms, i.e. they are seen only in one of the three copies, and the rest (130) are shared, i.e. seen in at least two copies. We observe a striking difference in the number of shared polymorphisms between long introns on one side and short introns and coding sequence on the other side. In long introns only 7% of the polymorphisms are shared, and they are only shared between the two more recent duplicates. In contrast, in short introns 78% and in CDS 67% of the polymorphisms are shared. A second consequence of IGC between genes is a generally higher density of polymorphic sites in short introns than in long introns (0.114 per bp vs 0.042 per bp; 2-sample test for equality of proportions with continuity correction: p = 8.6 *×* 10*^−^*^12^).

**Figure 3:**
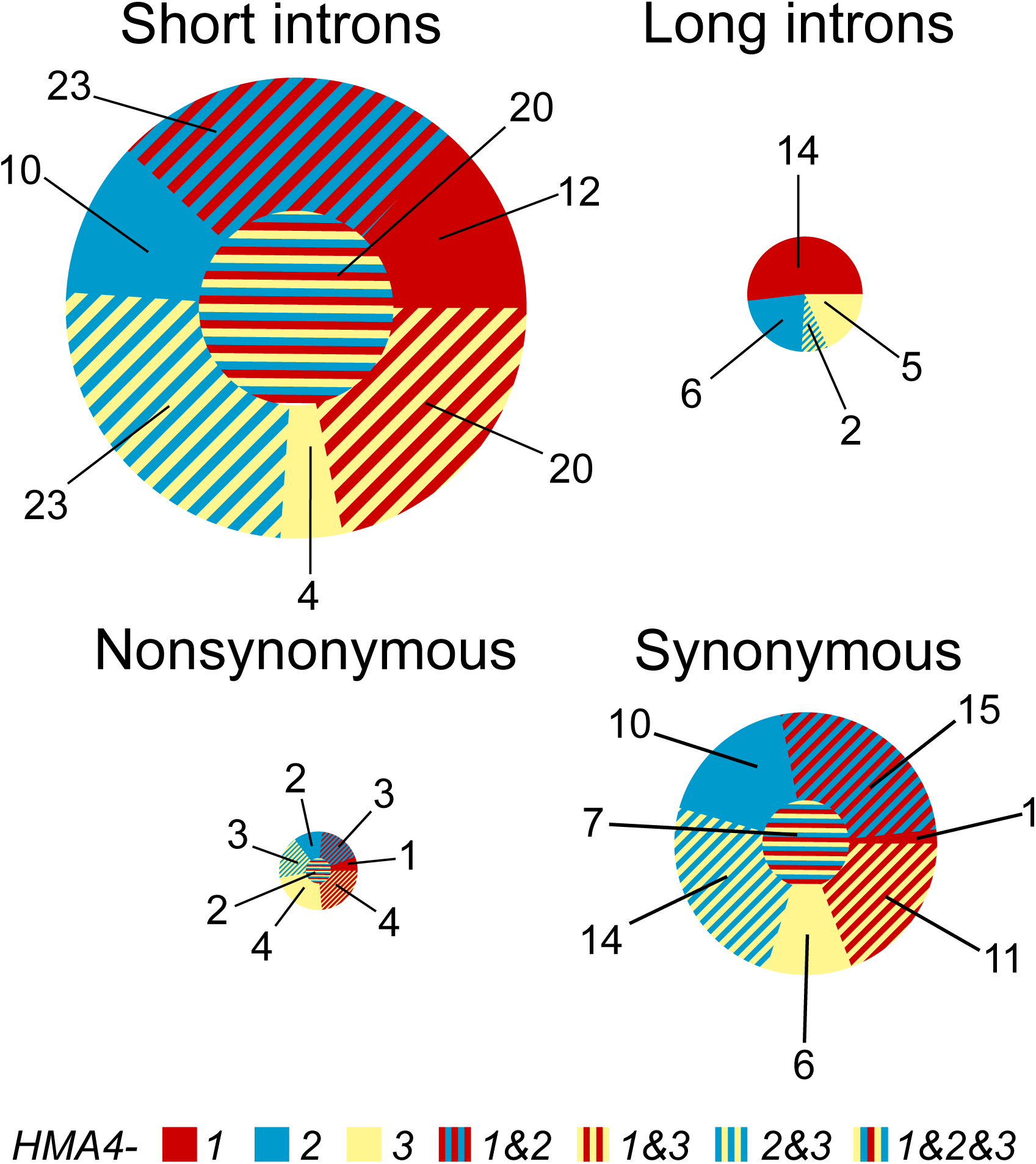
Pie charts depicting the numbers of variable sites per kilobase (SNP densities) in different parts of the *HMA4* genes. Variable sites are categorized as private to one copy (one color), shared between two (striped, two colors, in the outer ring), or shared among all three copies (striped, three colors, in the inner circle). Within a pie chart, all areas are proportional to the frequencies of variable sites private to a gene copy or shared between gene copies. Pie charts are proportional to each other based on their overall frequencies of variable sites.

**Table 2:** Total counts (and percentages) of shared and private polymorphisms in *AhHMA4* genes.

| *HMA4*- | length* | private polymorphisms |  |  | shared polymorphisms |  |  |  | total |
| --- | --- | --- | --- | --- | --- | --- | --- | --- | --- |
|  |  | 1 | 2 | 3 | 1&2 | 1&3 | 2&3 | 1&2&3 |  |
| intr<sup>short</sup> | 647 | 8 (11) | 7 (9) | 3 (4) | 15 (20) | 13 (17) | 15 (20) | 13 (17) | 74 |
| intr<sup>long</sup> | 2276 | 51 (53) | 25 (26) | 20 (20) | 0 | 0 | 7 (7) | 0 | 96 |
| CDS<sup>syn</sup> | 1161 | 1 (1) | 8 (15) | 5 (9) | 12 (23) | 9 (17) | 11 (21) | 6 (11) | 52 |
| CDS<sup>nonsyn</sup> | 2321 | 5 (8) | 6 (10) | 11 (18) | 10 (17) | 11 (18) | 9 (15) | 6 (10) | 58 |

\*average aligned sequence length in bp without alignment gaps

This suggests that long introns escape IGC. Therefore, their pairwise divergence should be a suitable measure to estimate the duplication times. Assuming a mutation rate of 7 *×* 10*^−^*^9^ per site per generation (Ossowski et al., 2010) and an effective population size of 83,000 (Roux et al., 2011), the observed pairwise differences suggest that the first duplication occurred approximately 4.34 (when comparing *HMA4-1* with *HMA4-2*) or 3.27 (when comparing *HMA4-1* with *HMA4-3*) million generations ago. Comparison of *HMA4-2* and *HMA4-3* with the same assumptions, suggests that the second duplication occurred around 166 thousand generations ago. Assuming a generation time of two years (Roux et al., 2011), the duplication events would date to approximately 7.6 (averages of 8.7 and 6.5) mya and 332 kya. The number calculated for the first duplication differs substantially – roughly by a factor of 20 – from the estimate of 357 kya derived by Roux et al. (2011) while our estimate for the second duplication is close to their estimate of 250 kya. One factor contributing to the large discrepancy is IGC: when neglected although present, time is severely underestimated. Furthermore, the presence of additional elements, e.g. repeats such as SINEs and Tc1/mariner in the long introns of the *HMA4* genes, may also contribute to imprecise estimates of divergence time.

Upon closer examination of the genetic variation in the coding sequences, we observed an enrichment of fixed differences at synonymous and nonsynonymous positions between *HMA4-1* and the two other paralogues in exons I and II and at the start of exon III (Figure 4, Supplementary table S1). These exons are separated from each other by long introns. Exon I and the start of exon II encode the *N* -terminal, cytosolic part of the protein. In the region encoding the transmembrane helices (exons II to IXa), synonymous variation exceeds nonsynonymous variation in all gene copies, indicating purifying selection conserving the amino acid sequence. However, the pattern observed in the cytosolic *C*-terminus (CC) is opposite in all three genes: there is more nonsynonymous than synonymous variation. Whether this is just due to relaxation of selective constraint or to positively selected, adaptive differences is a topic for further investigation.

**Figure 4:**
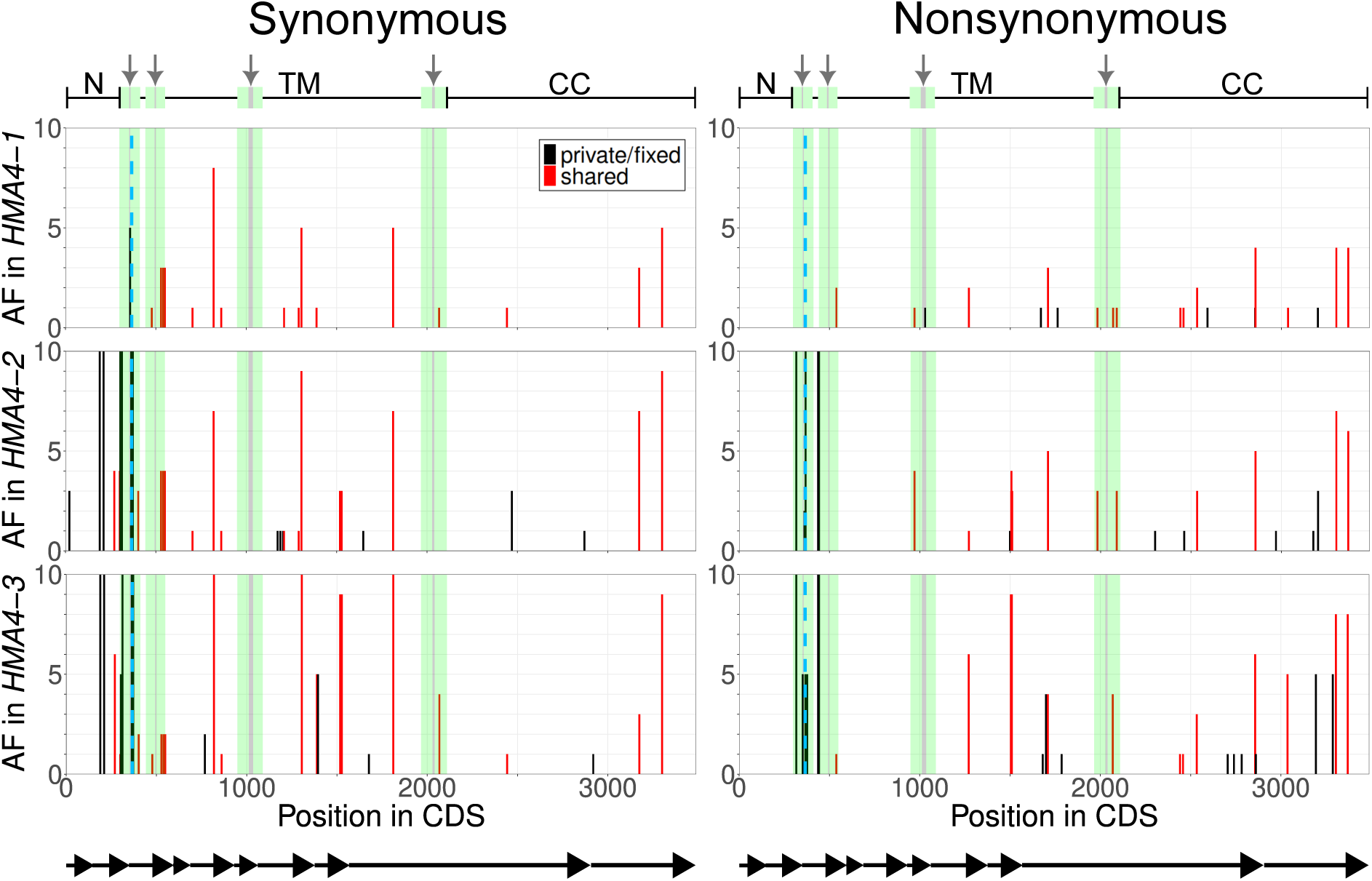
Absolute allele frequencies (AF) of variable positions in the coding sequences of the *HMA4* genes, split by the effect of the variant on the amino acid sequence. In each position, the allele found in the *HMA4-1* gene of accession Lan3.1 is taken as the reference. The plotted allele frequencies are those of all alternative alleles added together. Segments encoding transmembrane helices are marked in green, segments encoding amino acids which face extracellular are marked by grey vertical lines and highlighted by grey arrows. A blue, dashed line separates the coding exons I and II from the coding exons III-IX. Below the plots, a scheme depicts the exon boundaries. Above the plots, a scheme depicts the underlying structure of the protein, including the cytosolic N-terminus (N), the transmembrane domain region (TM) and the cytosolic regulatory C-terminus (CC).

### Modeling gene duplication and IGC

#### Two copies without IGC

To investigate the potential influence of IGC and different modes of selection on sequence evolution at the *HMA4* locus and similarly evolving loci, we developed a forward-in-time simulation model and adapted it based on our findings at this locus. We start with a diploid, panmictic population of effective size *N_e_* = 2000 in mutation-drift equilibrium. Evolution proceeds in discrete time and with non-overlapping generations. Initially, all individuals have a single copy of the gene of interest of size *L* = 20 kb. In generation 1, a duplication occurs in a single haplotype, creating a copy adjacent to the first gene. The new copy provides an adaptive benefit to the individual, and fitness is additive: homozygotes with one copy have a fitness of 1, heterozygotes with one extra copy have fitness 1 + *s_a_* and homozygotes with two copies in both haplotypes have fitness (1 + *s_a_*)^2^ *≈* 1 + 2*s_a_*. In our simulations, we used *s_a_* = 0.01. Reciprocal recombination (standard meiotic recombination) between the two loci occurs at a rate of *r*_rec_ per gamete per generation, while recombination within copies is neglected. We let the population evolve over 100 k generations and measured genetic diversity (*θ_π_* and *θ_w_*) within each copy, their ratio *θ_π_/θ_w_* and divergence between copies. As expected, divergence increases linearly with time, reciprocal recombination has almost no effect (Figures 5A and S2). The fixation process of the second, beneficial gene copy disturbs genetic diversity (here *θ_π_*) at the first locus when recombination is low (Figure 5C). This is easily explained by the hitchhiking effect which causes a reduction of genetic diversity at nearby loci. Shortly after fixation (at about *t* = 15 k generations), genetic diversity settles at the mutation-drift equilibrium so that the rates of mutation and loss or fixation due to genetic drift are balanced. This effect is also reflected in the ratio of the two quantities *θ_π_* and *θ_w_* (Figure 5E). When both loci are analyzed together, this hitchhiking signal can even be seen under high recombination (Figure 5G). While within-copy diversity stagnates after fixation (Figure 5C), the ratio continues to increase beyond 1 due to the accumulation of divergence between the two copies.

**Figure 5:**
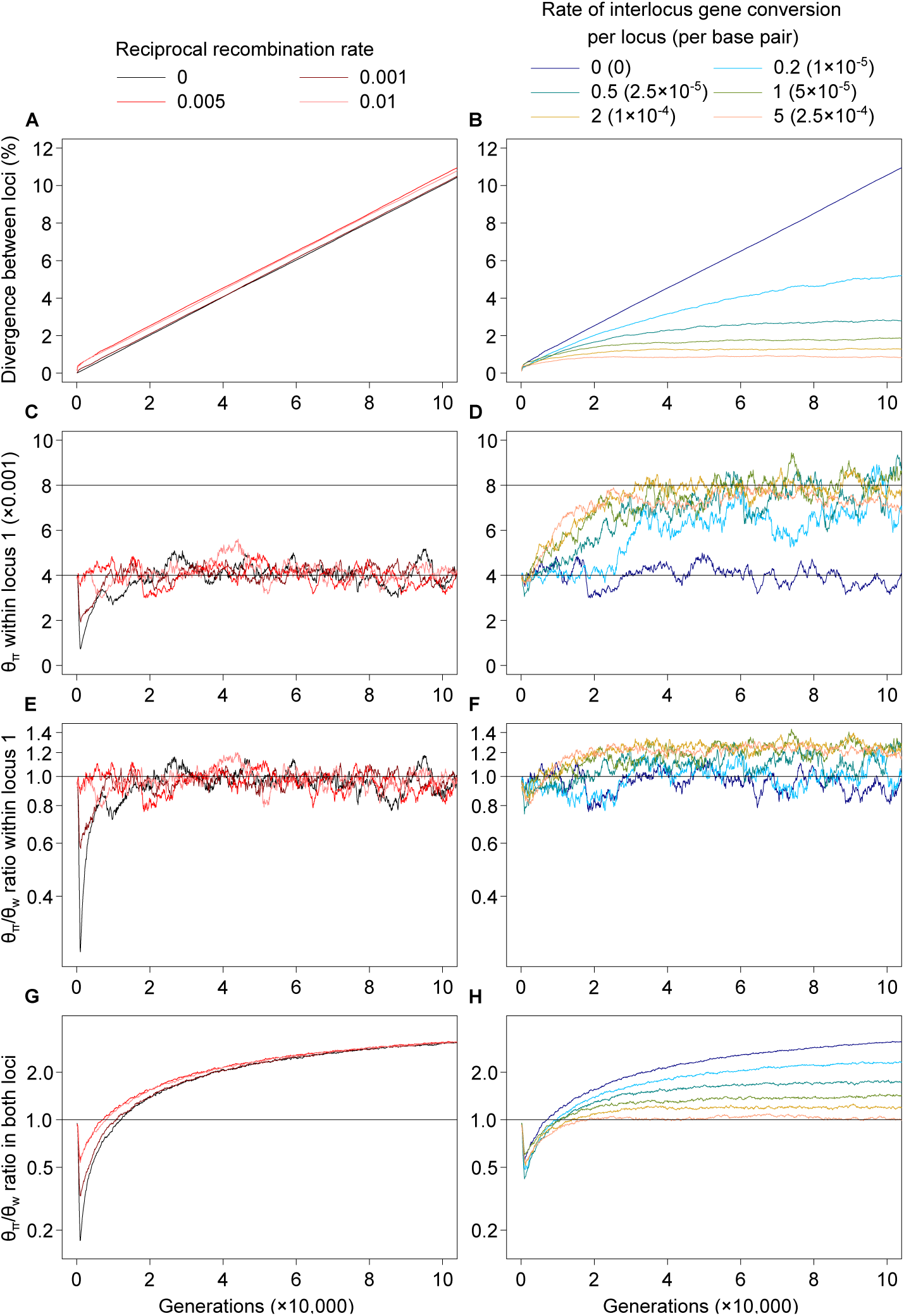
Variation statistics for a two-locus neutral model without and with IGC as a function of time. Reciprocal recombination rate between loci is *r*_rec_ = 0, 0.001, 0.005 or 0.01 (without IGC; panels A,C,E,G) or kept constant at *r*_rec_ = 0.005 (with IGC; panels B,D,F,H). A,B: mean sequence divergence between loci 1 and 2 since time of duplication *x* generations ago. C,D: pairwise nucleotide diversity (*θ_π_*) within locus 1. The horizontal line at *θ_π_* = 0.004 marks the single-locus expectation at mutation-drift equilibrium. Locus 2 dynamics differ from locus 1 only in the first few thousand generations (see Figure S2). E,F: *θ_π_/θ_w_* ratio for locus 1, expected to be 1 (horizontal line) in an equilibrium population; a ratio greater (smaller) than 1 is equivalent to a positive (negative) Tajima’s *D* statistic. G,H: *θ_π_/θ_w_* ratio for pooled data from both loci.

#### Two copies with IGC

IGC is the non-reciprocal transfer of a piece of homologous DNA from one gene to another. We assume that the length of the transferred segment, the *tract length L*_GC_, is on the order of some ten to a few hundred bp and much smaller than the gene length *L*. Let *L*_GC_ be a geometrically distributed random variable with mean 500 bp (Mansai et al. (2011)) and assume that IGC events occur at a rate of *r*_GC_ per generation per haplotype per (donor-to-target) direction. Then, 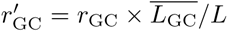 is the probability that a given base pair experiences IGC per direction per haplotype per generation. The per-locus IGC rate is 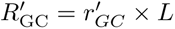, the expected number of base pairs affected by IGC per direction per haplotype per generation across the entire locus. We use the latter as a parameter in the simulations with values 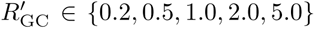. In the simulations with IGC we keep the reciprocal recombination rate fixed at *r*_rec_ = 0.005.

Repeated events of IGC homogenize the affected sequences, thereby suppressing the accumulation of divergence between the gene copies (Figure 5B). Loci experiencing IGC reach an IGC-mutation equilibrium, where the divergence between loci is stabilized at around *µ/*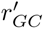, dependent on conversion rate, mean tract length and mutation rate. This equilibrium balances mutation increasing sequence divergence at rate *µ* and IGC reducing it at rate 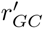. Notice that population size cancels out in this ratio. As a result, if the age of a gene duplication is calculated based on a molecular clock assuming neutrality, the duplication time will be overestimated shortly after the duplication due to reciprocal recombination increasing divergence, but severely underestimated later on due to IGC.

While it suppresses divergence, IGC raises genetic diversity within the loci (Figures 5D and S2). Depending on the rate of IGC, diversity may increase up to a factor 2 (in the case of two loci) as new mutations are transferred between the loci. Note that not only *θ_π_* but also *θ_w_* is affected by IGC. However, they are not equally affected: when the IGC rate is high, the relative increase of *θ_π_* is faster than that of *θ_w_*, indicated by their ratio being (slightly) above 1 (Figure 5F). Finally, when diversity of both loci is considered jointly, the time trajectory of the *θ_π_/θ_w_* ratio shows a dependence on the IGC rate (Figure 5H): at its highest, IGC can keep the *θ*-ratio around 1. However, the reduction of the ratio below 1 in the early phase until fixation of the new copy is unaffected.

#### Adding a third locus

Next, we consider a three copy model with two temporally separated duplication events to better match the application to the *HMA4* cluster in *A. halleri*. The three copy simulations start with a first duplication event at generation 1, as described before. At generation *T* = 10, 000, a haplotype with two loci is chosen and a third copy is generated by duplication of copy 2 on this haplotype. As before, we assume that the third copy confers an additive benefit of *s_a_* to the existing fitness of an individual (i.e., 2*s_a_*, when homozygous). As seen before, divergence continues to scale linearly with time in the absence of IGC. Thus, the two younger duplicates (loci 2 and 3) are diverged less from each other than either of them is from the original locus at all times (Figure 6A). Even if IGC is rare (*R*_GC_ = 0.2), this difference is erased quickly, within less than 10k generations (Figure 6B). Therefore, IGC may not only impede the inference of the timing of duplication events, but also the inference of the correct branching order.

**Figure 6:**
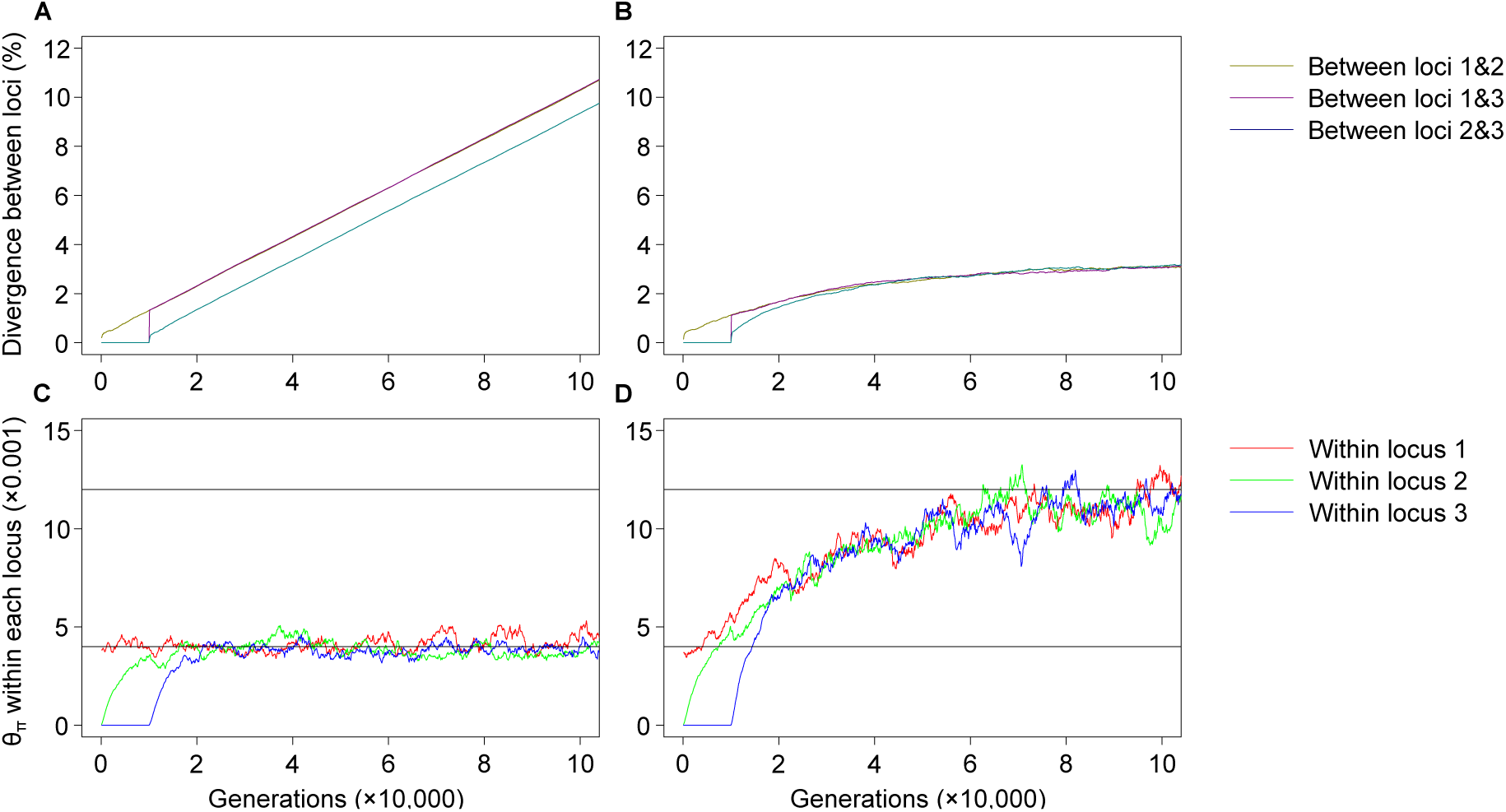
Within-locus diversity (A, B) and between-locus divergence (C, D) in a three locus neutral model without IGC (A, C) and with IGC (B, D; *R*_GC_ = 0.5).

#### Considering exon/intron structure of genes and variable IGC rates

Functional constraints differ across coding and non-coding regions and shape sequence divergence between genes. This in turn affects the rates of IGC which requires donor and target sequences to be similar. To examine the interplay of functional constraints and IGC, we incorporated the action of purifying selection and variable rates of IGC into the two locus model. A locus of *L* = 20 kb is divided into neutral introns and functionally constrained exons. We adopt a codon structure and treat every third codon position as neutral and the first two positions as subject to purifying selection (“nonsynonymous” sites). Mutations at nonsynonymous sites confer a multiplicative fitness effect of 1 *− s_p_* per mutated site. Simulations start from the neutral equilibrium and purifying selection on nonsynonymous sites is applied for a burn-in period of 40, 000 generations before the first duplication event occurs at generation 1. The negative fitness effects act independently of the beneficial selection on copy number. Both are multiplied to calculate an individual’s fitness (1 + *s_a_*)*^C−^*^2^ *×* (1 *− s_p_*)*^m^*, where *C ≥* 2 is the number of copies and *m ≥* 0 is the number of mutations on nonsynonymous sites, on both haplotypes. In addition to the functional constraint, we introduce a rate of IGC that is dependent on divergence (Teshima and Innan, 2004; Thornton, 2007): an IGC event still initiates with rate *R*_GC_, but it successfully terminates with probability *ϵ*, which decreases exponentially with increasing sequence divergence between donor and target sites within the IGC tract. Thus, the actual rate of IGC is reduced by the factor *ϵ* = *e^−dλ^*, where *d* is the percent sequence divergence within the conversion tract and *λ* is a ‘penalty’ parameter determining the strength of the reduction.

Due to the combined effect of purifying selection and IGC, nonsynonymous sites accumulate little to no sequence divergence over a range of parameter values of *R*_GC_ and *λ* (Figure 7). In contrast, synonymous sites and introns are both not exposed to purifying selection and accumulate more mutations than nonsynonymous sites, which may eventually contribute to divergence. However, synonymous sites are closely linked to nonsynonymous sites. Therefore, they are more likely covered by IGC tracts containing many nonsynonymous sites. As a result, synonymous sites diverge more slowly than introns. The combined effect is a mosaic pattern of the level of divergence, from nonsynonymous to synonymous sites to introns (Figure 7). For some parameter combinations (e.g. *R*_GC_ = 0.2, *s_p_* = 0.001), this can lead to a situation where divergence reaches an equilibrium in synonymous sites while introns continue to diverge for a longer time before reaching an equilibrium or without reaching an equilibrium (Figure 7A,B,D). Only when the IGC rate is sufficiently high and penalty is low, introns also quickly enter an IGC-mutation equilibrium in which divergence ceases to accumulate (Figure 7C). This is consistent with the observation from the *HMA4* genes that the divergence in (long) introns between gene copies can exceed the nucleotide diversity found within gene copies while synonymous sites are highly similar (Figure 2).

**Figure 7:**
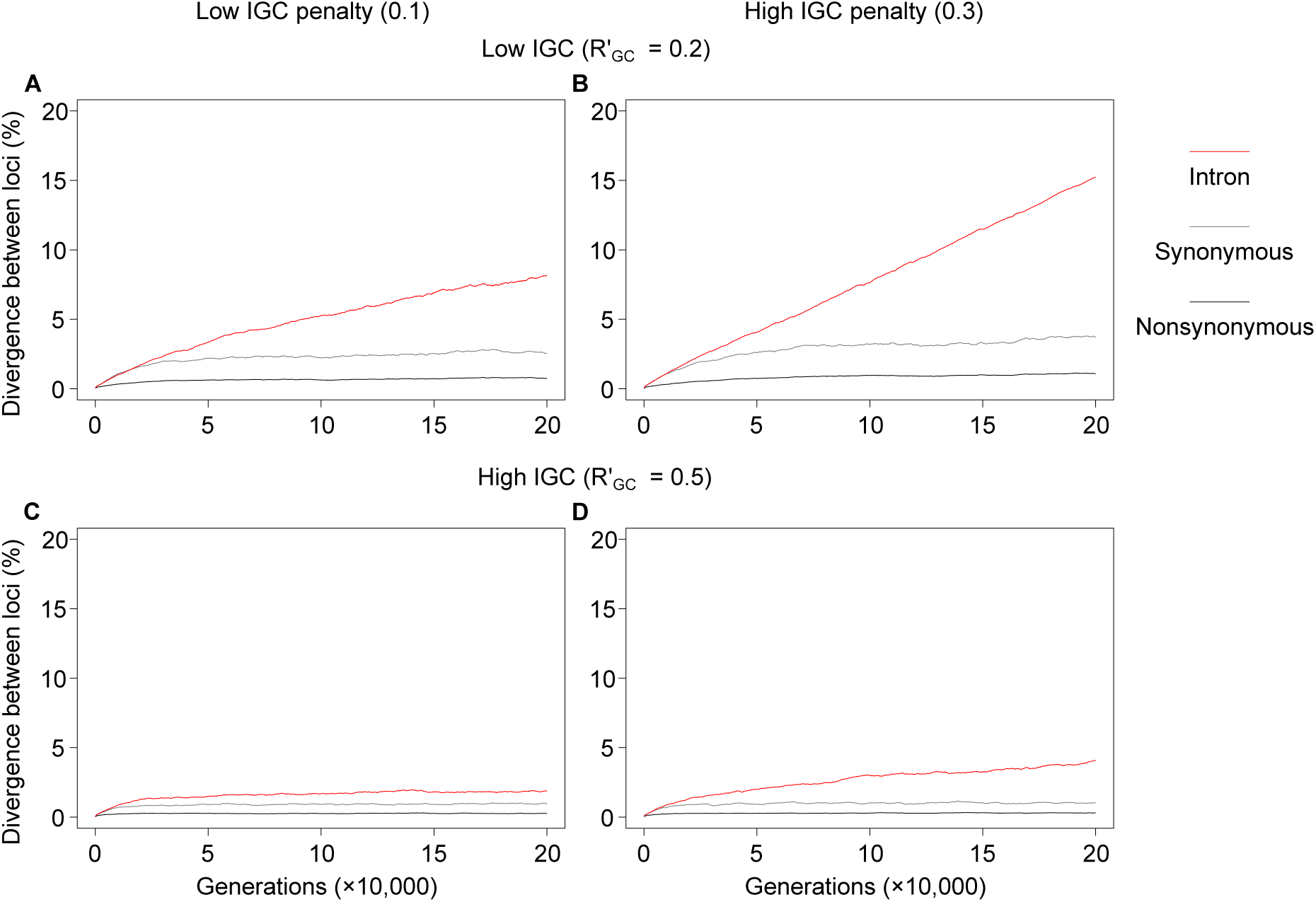
Mean sequence divergence between two loci evolving under selective constraint and a rate of IGC dependent on divergence. The sequence is divided into nonsynonymous positions (black), synonymous positions (grey) and introns (red). Columns: penalty *λ* is low (0.1; A, C) or high (0.3; B, D). Rows: IGC is low (*R*_GC_ = 0.2; A, B) or high (*R*_GC_ = 1.0; C, D). Purifying selection is weak (*s_p_* = 0.001).

#### Considering variable intron sizes and IGC tract lengths

We observed that not all introns of the *HMA4* locus are equally affected by IGC. We suspect that this may be related to different intron lengths. To investigate this possibility, we modeled a locus of 20 kb with 6 introns of sizes ranging from 60 bp to 3 kb. One intron of size 2840 bp, here called “terminal” intron, is flanked by only one coding exon and can be thought of as a 3’ UTR. We also tested the effect of varying IGC tract length. We assumed that tract lengths are chosen from a geometric distribution with mean 100, 500 or 1000 bp) and ran the simulations for 200, 000 generations.

We found that sequence divergence increases with intron size: while shorter introns diverged at a similar rate as synonymous positions, longer introns, and especially the terminal intron, diverged much faster (Figure 8). The tract length of IGC also affects sequence divergence: shorter tract lengths allow introns to escape the linked effects of purifying selection operating on nonsynonymous sites in adjacent exons, thus enabling greater divergence (Figure 8A,D). As tract size increases, the accumulation of divergence in introns decreases (Figure 8B,C,F). The greatest differences in divergence between long and short introns are observed when the mean tract length is intermediate and IGC is under high divergence-dependent penalty (Figure 8E). The differences between long and short introns observed in the *HMA4* genes of *A. halleri* may be driven by conserved, IGC-affected exons on adjacent intronic regions.

**Figure 8:**
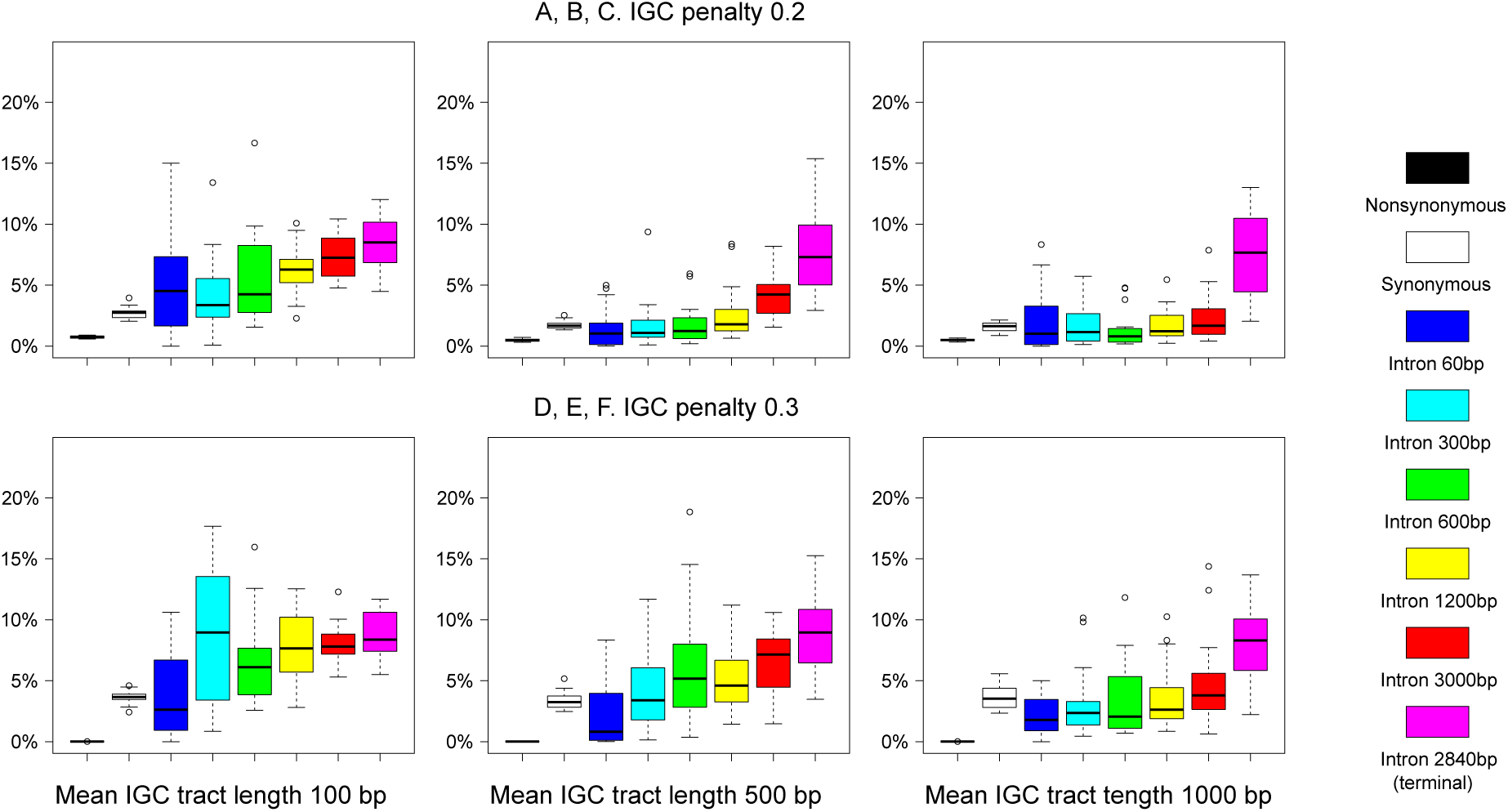
Pairwise sequence divergence between copies at generation 200, 000, given different values of *s_p_* = 0.001 (strength of purifying selection), *λ* = 0.2 or 0.3 (IGC penalty) and different mean IGC tract lengths. The IGC rate *R*_GC_ = 0.5 is kept constant. The ‘terminal’ intron is flanked by a coding exon only on its 5’ side.

#### Evolution of the AhHMA4 locus in the light of the model

Finally, we adapted the number and lengths of coding exons and introns in the model to those in the *HMA4* genes, from the 5’ UTR to the 3’ UTR. Since intron lengths varied across the empirical data, we used average lengths. We modeled a single duplication event only, since a second event does not introduce additional qualitative insights. During the simulations, we recorded the ratio of shared to non-shared polymorphisms over time (Figure 9A). After around 100k generations, around 18-20% of polymorphisms in synonymous sites and short introns are shared among gene copies. In contrast, only 5% of polymorphisms in nonsynonymous sites are shared, presumably because purifying selection (*s_p_* = 0.001) is acting on both gene copies. As observed in the previous model scenario, long introns diverge with time and escape IGC. As a consequence, polymorphisms which were previously shared between genes become fixed for one or the other allele and fewer new polymorphisms are transferred between loci. Short introns and protein-coding exons on the other hand continuously experience IGC and share polymorphisms between genes. These results broadly reflect the empirical observations in the *HMA4* genes, except that the simulated genes show a lower number of shared polymorphisms at nonsynonymous sites (Figure 3). One reason for this discrepancy could be the model assumption of an identical effect of purifying selection on all nonsynonymous sites. In reality, there is a plausible possibility that selective pressure varies across different regions of the *HMA4* locus (see Figures 4 & S1).

**Figure 9:**
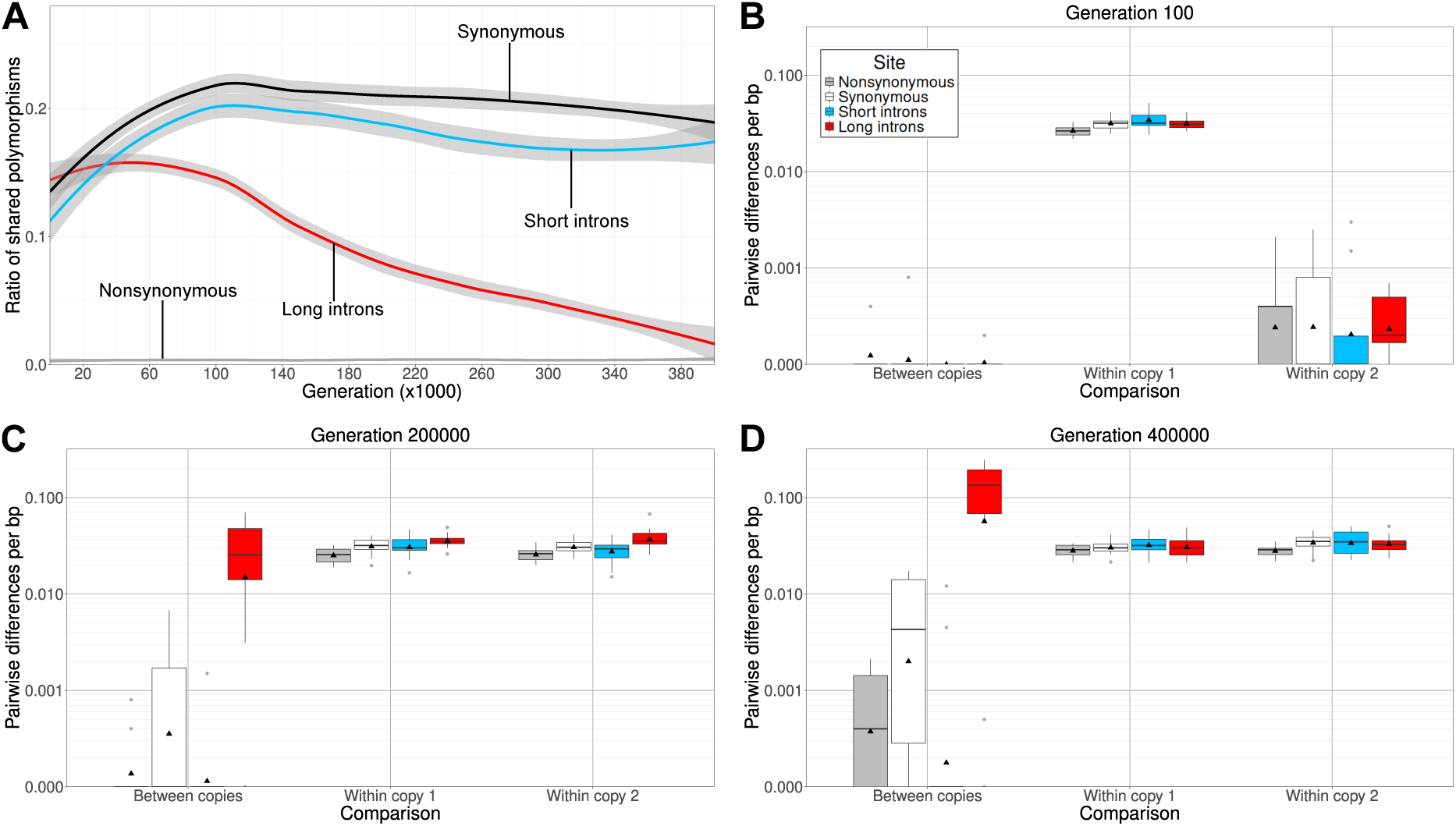
Results from the simulations with a gene model adapted to the gene structure of the *HMA4* genes of *A. halleri*. A: Ratio of polymorphic sites which were shared between both simulated genes. The remaining polymorphisms only occurred in either one of the two gene copies. B-D: Pairwise differences per bp within a gene copy and between the two gene copies, split by long introns, short introns, synonymous and nonsynonymous sites. For all shown results, the rate of GC (*R*_GC_ = 0.7), purifying selection (*s_p_* = 0.01) and IGC penalty were high (*λ* = 0.5).

Furthermore, at intervals of 100 generations we took snapshots of the population and recorded additional quantities. Shortly after duplication, the new copy shows little divergence. Allelic variation is present only in the ancestral gene (Figure 9B). After 200k generations, genetic diversity at the new copy has reached the level of the ancestral gene (Figure 9C) at the mutation-drift equilibrium (at about *θ_π_* = 0.04). Between genes, synonymous sites have diverged a little and long introns have diverged to a level exceeding within-gene diversity. After another 200k generations, long introns have diverged even further (Figure 9D) while synonymous sites and short introns are starting to approach the level of within-gene diversity. Therefore, after 200k generations, shared polymorphisms, divergence between long introns and within-gene diversity reflect the observations in the *HMA4* genes while divergence between IGC-affected regions lags behind.

## Discussion

We revisited the *HMA4* gene copies in *Arabidopsis halleri* and reconsidered its evolutionary history to design and refine a simulation model of gene duplication and interlocus gene conversion. The model generalizes earlier models (e.g. by Teshima and Innan (2004); Thornton (2007); Hartasánchez et al. (2014)) by incorporating not only IGC but also reciprocal recombination and natural selection as evolutionary forces and by considering a realistic exon/intron structure of the modeled genes. One of our goals was to reassess the age estimates of the two duplication events which led to copies *HMA4-2* and *HMA4-3*. The exon/intron structure of the three genes is highly conserved, suggesting strong functional constraint, recent duplication events or both. The HMA4 protein is encoded by ten exons; it consists of a short cytosolic *N* -terminus, a transmembrane domain with eight TM helices and a large unstructured *C*-terminal cytosolic domain, which was proposed to have a regulatory role (Hanikenne et al., 2013). Across a sample of ten accessions, we observe an uneven distribution of shared *vs.* non-shared and of synonymous *vs.* nonsynonymous polymorphisms along the genes. For instance, shared, synonymous polymorphisms are mostly clustered in the TM domain and nonsynonymous polymorphisms are strikingly overrepresented in the *C*-terminal domain. Since patterns of genetic diversity and levels of divergence are far from uniform and do not exhibit any clock-like behavior, it is clear that different evolutionary forces are at work in different parts of the genes. In agreement with this, there is also far more divergence from the *A. lyrata* and *A. thaliana HMA4* genes in the C-terminal cytosolic domain than in the transmembrane domain.

With our computer simulations, we tracked diversity and divergence forward in time beginning with a hypothetical duplication event which occurred in one individual leading to a new copy with a fitness benefit and then quickly became fixed in the population. Genetic diversity at the original locus is initially reduced as a result of hitchhiking, i.e., of positive selection and tight linkage, but shortly after fixation it increases - due to IGC between loci - to twice of its original mutation-drift equilibrium level. In the long run, IGC generates signatures in the site frequency spectrum that resemble balancing or diversifying selection, which could lead to misinterpretation in empirical datasets Hanikenne et al. (2013). Concurrently, IGC erodes signals of the molecular clock by homogenizing sequences between genes and retarding – more or less severely – the accumulation of sequence divergence compared to a scenario without IGC. This makes the determination of the timing and order of gene duplication events difficult, if not obsolete.

To more accurately capture the complexity of real gene families, such as *AhHMA4*, we incorporated exon/intron structure of genes and a distinction between synonymous and nonsynonymous sites into our model. These adjustments revealed an intricate interplay between IGC and natural selection. For example, as duplicated genes accumulate new mutations, their increasing sequence divergence can reduce or even eliminate the potential for IGC. However, in regions under strong purifying selection, especially in coding regions, sequence similarity is maintained, enabling prolonged IGC activity. This persistence of gene conversion generates a pattern of concerted evolution between copies, masks their evolutionary independence and impedes divergence. Within a gene, certain gene parts may be affected by recurrent IGC and trapped in a state of concerted evolution, while other parts – typically those which are under no or only little selective pressure – escape IGC and accumulate differences. As a result, the characteristic signatures of IGC (reduced divergence between copies and excess of shared polymorphisms) are only observable in conserved regions and those close to them. The patterns emerging from our simulations are consistent with these empirical findings and align closely with our observations in the *HMA4* gene family. Specifically, long introns appear to escape IGC in the long run, while coding exons and short introns remain subject to its influence. This is reflected by the observed divergence between long introns of *AhHMA4-1* and those of the more recent duplicates which exceeds within-copy diversity, and by the lack of polymorphisms shared between these long introns.

Nevertheless, certain observations from the *HMA4* paralogues were not fully captured in our model. For instance, the model predicts a higher proportion of shared polymorphisms at synonymous sites than at nonsynonymous sites. In contrast, the empirical data from *HMA4* genes reveal that shared polymorphisms were equally abundant at these two types of sites (see Figure 3). This discrepancy may arise because the model assumes uniformly strong purifying selection across all nonsynonymous sites, whereas only parts of the *HMA4* coding sequence may be under positive selection (Clarke, 1970). Alternatively, purifying selection may be weaker in reality than assumed in our simulations. Perhaps only a small fraction of nonsynonymous variants shared between *HMA4* genes is affected by (strong) purifying selection. In fact, further analysis of these shared polymorphisms shows that they often result in substitutions with biochemically similar amino acids, e.g., exchanges of different hydrophobic amino acids (see supplementary data file S1).

Moreover, while empirical variation patterns differ among exons, the model does not recover such differences. For example, from the sequences of the *HMA4* genes, we observed fixed differences between *HMA4-1* and the other two copies and these differences were restricted to the coding exons I and II. In contrast, the other exons showed more shared polymorphisms and no fixed differences. It is possible that the close proximity of these two exons to the long introns affect IGC across the entire region. This may facilitate further sequence differentiation between gene copies and promote independent evolutionary trajectories. The overall gene architecture (e.g. intron and exon number, positions and length) is shared to a large extent with other closely related species, *A. lyrata* and *A. thaliana* (Figure 1), indicating that this gene architecture is an ancestral feature. It may directly impact the likelihood of IGC and have consequences on how sequence divergence is affected by neutral and selective processes.

Another striking observation is the distribution of nonsynonymous variation across the genes. The cytosolic, regulatory *C*-terminus exhibits an excess of nonsynonymous variation in contrast to the *N* -terminus. One possible explanation is that the *C*-terminal domain is less constrained than the transmembrane domain. Although it contains 11 di-cysteine motifs which were proposed to contribute to the binding of Zn^2+^ ions and are essential for the function of the protein (Lekeux et al., 2018), other parts of the *C*-terminus might even be subject to diversifying or positive selection, which can be promoted by IGC (Mano and Innan, 2008). AlphaFold (Jumper et al., 2021) predicts that the *C*-terminus is unfolded (entries Q2I7E8 for *A. halleri* and O64474 for *A. thaliana*), but has low confidence in its prediction. It is possible that the HMA4 proteins have an intrinsically disordered C-terminal domain (Ruff and Pappu, 2021) so that the di-Cys residues are accessible and the protein conformation can change upon binding (Porter and Looger, 2018). However, it is also possible that the fold could not be predicted from the amino acid sequence because it is structured in a way unknown to AlphaFold or because its proper function may require post-translational modifications (Burén et al., 2011), chaperones (Gething and Sambrook, 1992) or other co-factors. The exact mechanisms of ion binding currently remain unclear Lekeux et al. (2018).

Although IGC does not directly impact the introduction of base substitutions and amino acid changes, it parallelizes their accumulation among adjacent gene copies (Yang et al., 2023). It therefore may prevent neofunctionalization and subfunctionalization in multi-gene families (”concerted evolution”). If functional differentiation is favored by natural selection, mechanisms that reduce IGC may be beneficial. Such mechanisms could include translocations or inversions of gene copies or the acquisition of large insertions in or near gene copies. Although IGC can occur between non-adjacent genes and even between loci on different chromosomes, its frequency decreases with genomic distance (Xu et al., 2008). Conversely, close physical proximity and high sequence similarity between duplicated loci does not necessarily guarantee the occurrence of IGC. Several other factors influence the likelihood and frequency of IGC, such as local recombination rate (Henderson, 2012), GC-content (Muyle et al., 2011), genomic stability (Taghian and Nickoloff, 1997), DNA methylation (Yelina et al., 2015) and transcriptional activity (González-Barrera et al., 2002). These factors can either facilitate or suppress gene conversion, depending on the genomic context. Generally, IGC is likely more widespread than traditionally recognized (e.g. Yang et al. (2023)). While there are prominent and well-known examples of IGC-induced concerted evolution (e.g., the histone H1 gene family (Eirín-López et al., 2004) or immune-related gene families (Zid and Drouin, 2013)), new findings are being reported (e.g., partial gene conversion affecting transposable elements (Doronina et al., 2021) or gene families (Shen et al., 2021)). Anyway, the presence and potential impact of IGC deserves more attention in genomic studies of gene family evolution.

Returning to our goal of dating duplication events in *AhHMA4*, we draw the following conclusions: we expect that previous estimates of the time of the first duplication (357 kyrs ago, Roux et al. (2011)) have underestimated the impact of IGC and placed the estimate too close to the present. Such a recent duplication time also seems unlikely given that the three copies show a very similar level of genetic diversity in both short and long introns (Figure 2), suggesting that mutation-drift equilibrium has been reached in all gene copies. The time to reach this equilibrium under neutrality scales with the inverse of the mutation rate – *O*(1*/µ*) = 100 million generations – which is vastly different from the derived estimate. On the other end, an upper bound to the age estimate seems to be defined by the speciation time from closely related species in the genus *Arabidopsis* since neither *A. thaliana* nor the closely related *A. lyrata* have a triplication of *HMA4*. It is a single copy gene in *A. thaliana* and two gene copies, however on different chromosomes, are annotated in *A. lyrata*. This suggests that both duplication events, and at least the younger one, occurred after the split of *A. lyrata* and *A. halleri*. Unfortunately, very incongruent speciation times, differing by a factor of almost 50, are reported for this split. They range from 0.3 mya (Roux et al., 2011) to 6.1 mya (Guo et al., 2021) and even 12.6 mya (Irisarri et al., 2021; Kumar et al., 2022). In any case, the *A. halleri* duplications are most likely not older than these upper bound extremes.

The separation of the sequence data that we undertook into exons, short and long introns revealed a very different divergence pattern in the long introns compared to the other sequence elements. It is consistent with the idea that the latter are homogenized among copies by ongoing IGC, while long introns escape IGC and carry a time-dependent signal of divergence. This idea is corroborated by our simulations. Without this divergence signal from the long introns, determining the duplication times as well as the branching order would be extremely difficult or even impossible. With the signal however, and guided by our model, we can confirm that *HMA4-2* and *HMA4-3* participated in the more recent duplication (either as the template and the other as the duplicate) and provide new estimates of the duplication times: we place the first duplication event at 7.6 mya and the second one at 0.3 mya.

Typically, long introns are difficult to align. For the age estimates we only considered SNPs in alignment blocks which could be aligned with high confidence. Indels were ignored and this may contribute to underestimating true sequence divergence and therefore to underestimating duplication time. Furthermore, after duplication, the entire gene should initially experience IGC, independent of intron length. Escape from IGC evolves only gradually, while mutations accumulate, except if it is facilitated via mutations affecting multiple bases at once such as inversions or TE insertions. This effect also biases the age estimates downward. Finally, generation time is a potential source of error. With *A. halleri* being a perennial plant, we assumed a generation time of 2 years (Roux et al., 2011). However, average generation time may be different or additionally vary among accessions and locations. If true, this would also shift our time estimates accordingly.

In summary, while the precision of age estimates of tandemly arrayed gene copies is impeded by IGC, our simulation- and data-informed estimates indicate that the first duplication event happened several million years ago, possibly contemporarily with the split from the closely related species *A. lyrata*, while the second duplication likely happened much more recently, just a few hundred thousand years ago. Therefore, in agreement with *HMA4* gene copy number expansion in *A. halleri* on both heavy metal-enriched and uncontaminated soils (Hanikenne et al., 2013), the ability of *A. halleri* to hyperaccumulate zinc without suffering from its toxicity did not evolve as a response to anthropogenic environmental change, but instead exists at least since the Miocene.

## Material and Methods

### Empirical data and sequence analysis

#### Extracting sequence segments corresponding to the HMA4 region

A contig comprising the complete *A. halleri HMA4* genomic region was assembled from the sequences of two BAC clones (EU382073.1 and EU382072.1, total length 289,748 nt; Hanikenne et al., 2008, 2013) using merger from the EMBOSS package (Rice et al., 2000). The sequence of this contig was used as a query in a stand-alone blast (v2.13.0; Camacho et al., 2009) against each of five chromosome-level genome assemblies of *A. halleri* : Lan3.1 v2.10 (Phytozome; *Arabidopsis halleri* v2.03, DOE-JGI), Pais 09 v1.1.0, Wall 10 v1.1.0, Lan5 Hap1 v1.1.0 and Lan5 Hap2 v1.1.0 (Stein et al., 2017). From the resulting alignments, the nucleotide sequence coordinates with the highest sequence similarities and coverages were used to extract the complete sequence segments of the *HMA4* genomic regions using samtools (v1.18; Danecek et al., 2020). To extract the *HMA4* genomic region from five further contig-level genome assemblies (Rund 05a v1.0.0, Goli 08 v1.0.0, Noss 08 v1.0.0, Bors 12 v1.0.0, and Ukan 25 v1.0.0) (Stein et al., 2017), we performed the following steps: we used the amino acid sequence of copy *AhHMA4-1* from the corresponding BAC sequence (GenBank ID EU382073.1; Hanikenne et al., 2008) to query the *Arabidopsis thaliana* protein database via TAIR (Huala et al., 2001). The best alignment was *At* HMA4 (Locus: AT2G19110). In the “topology” entry of Aramemnon (Schwacke and Flügge, 2018), we identified the highly conserved final 10 amino acids of the 8th transmembrane helix (TCLLVIFNSM) of *At* HMA4 according to ConPred v2 (Arai et al., 2004). This 10-aa sequence fragment was used as a query in a stand-alone blast (v2.13.0) to identify the nucleotide position of *Ah*HMA4-1 corresponding to the beginning of the cytosolic C-terminus. BAC positions 71,170 to 72,180 corresponding to the 704-aa cytosolic C-terminus of *AhHMA4-1* were used as a query against each of the contig-level assemblies using stand-alone blast (v2.13.0). For each assembly, we thus obtained the nucleotide sequence coordinates for the top three hits with more than 99 percent sequence similarity and 100 percent query coverage, which corresponded to the three tandem *AhHMA4* gene copies, all within a single contig. From each assembly, we then extracted the sequence fragment from 20,000 bp upstream of the first (smallest) nucleotide sequence position to 20,000 bp downstream of the last (largest) nucleotide sequence position using samtools (v1.18). Finally, the obtained sequence segments were confirmed to contain *AhHMA4-1*, *AhHMA4-2* and *AhHMA4-3* by using the nucleotide blast module in NCBI web against the core nucleotide database (core-nt; Camacho et al., 2009). The sequence information and accessions for all samples are available in the table supplementary data file S2. The *HMA4* tandem-duplicate sequences generated in this study have been deposited in GenBank under the accession numbers PZ740972-PZ741001 (30 sequences; three tandem copies - *HMA4-1*, *HMA4-2* and *HMA4-3* - for each of ten *A. halleri* accessions/haplotypes: the BAC 7C17 clone and the Lan3.1, Lan5 (Hap1, Hap2), Rund 05, Bors 12, Goli 08, Ukan 25, Pais 09 and Wall 10 accessions).

#### Annotation of HMA4 genomic regions

We used publicly available annotations of *HMA4* genes from the Lan3.1 genome (Lan3.1 v2.10; Phytozome; *Arabidopsis halleri* v2.1.0, DOE-JGI; Cheng et al., 2017; Rawat et al., 2015) to annotate promoter, 5’-UTR, 3’-UTR and intron-exon structure of the *HMA4* genes from all other *A. halleri* accessions (Castanedo et al., 2025). To facilitate this process, we conducted pairwise blast alignments for each previously annotated feature against each corresponding *HMA4* gene copy for each *A. halleri* accession. Graph-based representations were produced with the SnapGene software (SnapGene 4.2.11, San Diego, California).

#### Variant analysis

The *HMA4* genes were then aligned using T-Coffee and visually inspected. The annotations were manually checked for errors and inconsistencies by aligning all annotated exons and all annotated introns separately using T-Coffee followed by visual inspection. The long intron alignment was used to identify insertions of more than 30 bp which occurred in only one gene copy. The introns of all *HMA4* genes were separated into long introns (introns 2 and 3) and short introns (introns 4-10). The identified intronic insertions which only occurred in one gene copy were removed with sed. Coding sequences, short and long introns of all *HMA4* genes were separately aligned with T-Coffee. All alignments were checked for quality using the Transitive Consistency Score (TCS) from the T-Coffee web server (Chang et al., 2015). Long intron alignments were filtered column-wise with a score threshold of 6 to remove poorly aligned regions. Genetic distances in synonymous and nonsynonymous sites were calculated using a bash script. Genetic distances in intronic sites were quantified by creating a VCF file of all SNPs from the alignments via snp-sites (version 2.5.1; Page et al. (2016)) and calculating allele frequencies in all sites, in which neither of the compared pair of sequences contained a gap, using vcftools --freq (version 0.1.16; Danecek et al. (2011)). The numbers of variants which were private to one gene or shared between genes were quantified by extracting separate VCF files for each gene copy from the ones previously created, filtering for biallelic SNPs and calculating allele frequencies with vcftools --freq again. The polymorphic sites were subsequently visualized as pie charts in R. The positions of the transmembrane domains were predicted by running DeepTMHMM (Hallgren et al., 2022) on the amino acid sequence of the HMA4 protein from Lan3.1. All data analysis and visualization scripts are available at https://github.com/YSchaefer/HMA4_analysis.

### Model and computer simulations

All simulations are based on an in-house developed Wright-Fisher-type model written in R. These are the parameters of the model:

- *N_e_*: effective population size, assumed to remain constant at *N_e_* = 2000
- *L*: length of one gene in bp, assumed to remain constant at *L* = 20000
- *µ*_0_: mutation rate per bp per generation, assumed to remain constant at *µ*_0_ = 5*e*-7
- *µ_L_* = *µ*_0_ *L*: mutation rate per gene per generation
- *F_i_*(1 *≤ i ≤ L*): functional state of the *i*th site in a gene. *F_i_* = 1 if the *i*-th site is under selective constraint (nonsynonymous) and *F_i_* = 0 if the *i*-th site is neutral. If the *i*th site is in an intron, then *F_i_* = 0. If the *i*th site is in an exon, then *F_i_* = 0 if *i*%3 *≡* 0 (synonymous sites), and *F_i_* = 1 otherwise (nonsynonymous sites)
- *s_a_*: selective benefit of any additional gene copy in an individual. Selection is assumed to be multiplicative across alleles and across loci. At generation 0 all individuals have a single copy on both haplotypes and have fitness 1. Individuals with an additional copy have fitness 1 + *s_a_* if heterozygote, and (1 + *s_a_*)^2^ *≈* 1 + 2*s_a_* if homozygote (i.e., two copies on both haplotypes). In the three copy model maximum fitness is (1 + *s_a_*)^4^ *≈* 1 + 4*s_a_*, if an individual carries three copies on both haplotypes. Since we assume rapid fixation, all individuals quickly become homozygous for either two or three copies
- *s_p_*: selective cost of a mutation at a nonsynonymous site. Again, selection is multiplicative across sites
- Total fitness of an individual is *w* = (1 + *s_a_*)*^C−^*^2^ *×* (1 *− s_p_*)*^m^*, where *C* is the total number of gene copies on two haplotypes, and *m* is the total number of nonsynonymous mutated sites in each copy and each haplotype. At the start of the simulation (generation 0), *C* = 2 for all individuals and the population is assumed to be in equilibrium, produced by a burn-in over 40k generations. Mutations will already accumulate during burn-in, such that *m >* 0 at generation 0. Time proceeds in non-overlapping generations. Two individuals (parents) from the current generation are chosen independently, and with probability proportional to their fitness, to produce a child for the next generation. This procedure is repeated *N_e_* times so that population size remains constant
- *T_j_*: time (in generations) when the *j*th copy is created by duplication from a randomly selected haplotype with *j −* 1 copies. Here, *j* = 2 or 3 and *T*_2_ = 1.
- *r_j_*: reciprocal recombination rate (per individual per generation) between copy *j −* 1 and copy *j*. Here, *j* = 2 or 3.
- *r*_GC_: rate with which IGC events occur per copy per haplotype per generation (regardless of the tract length)
- *L*_GC_ is a geometrically distributed random variable representing gene conversion tract length. Mean tract length is 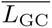 (maximum tract length is truncated at locus size *L*)
- 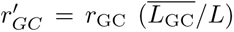: the rate of IGC per site per donor-to-target direction per haplotype per generation. That is to say, the probability that a given site experiences an IGC event from a given copy to another given copy in a given haplotype is *r^′^*
- *λ*: the factor by which the IGC rate is reduced as a function of sequence divergence. This is realized by first (randomly) choosing the IGC tract length and the IGC donor and target sites. Then, donor and target sequences are compared to obtain their divergence *D*. The initiated IGC event is completed with probability *ϵ* = *e^−Dλ^* and aborted with probability 1 *− ɛ*

Each simulation run starts with an equilibrium population with each haplotype having a single copy. Under neutrality, this initial state is generated directly with ms (Hudson, 2002). The initial population has an expected average nucleotide diversity of *θ* = 4*N_e_µ*_0_ = 4*e*-3 before application of selection. To account for selection, the ms-generated population is propagated for another 40k generations with the selection scheme explained above to (approximately) reach a selection-mutation equilibrium.

A generation in the simulation encompasses the following steps: (a) calculating fitness of each individual; (b) randomly choosing a pair of parents with random sampling weighted by fitness; (c) apply reciprocal recombination and combine gametes to form new individuals; (d) apply IGC (e) apply mutation on single nucleotide level.

Should a gene copy become lost before fixation, the population is returned to the state before generation of the copy by duplication, and re-established by duplication of a randomly chosen haplotype. Each run lasts 200, 000 generations, and population statistics are calculated every 100 generations. All simulation scripts are available at https://github.com/y-zheng/Interlocus_gene_conversion_modeling. Scripts for the simulations adapted to the exon/intron structure of the *HMA4* genes are available at https://github.com/YSchaefer/HMA4_analysis.

## Acknowledgments

This research was funded by the German Research Foundation (Deutsche Forschungsgemeinschaft, DFG) – Project-ID 456082119 – TRR 341/1, subprojects A9 and B5. The genome assemblies were funded by the European Research Council (ERC-AdG LEAP EXTREME grant 788380 to U.K.). We gratefully acknowledge the Genomics and Transcriptomics Laboratory of the Biological and Medical Research Centre (BMFZ) at Heinrich-Heine University (HHU) Düsseldorf, led by Prof. Dr. Karl Köhrer, where sequence data were generated. We thank Prof. Dr. Korbinian Schneeberger and Dr. Raul Wilfjes (LMU Munich, Germany) for the sequencing and the generation of two haplotype-phased genome assemblies for *A. halleri* accession Lan5.

## Declaration of competing interests

The authors declare no competing interests.

## Author contributions

YS: data and statistical analyses; contributed to sequence annotation; writing of the manuscript

YZ: mathematical modeling; data and statistical analyses; writing of the manuscript

AP, BP, PK: data analysis and sequence annotation VK: sequencing

UK: designed the project; supervision of sequencing, sequence annotation and data analysis; acquisition of funding; writing of the manuscript

LR, TW: designed the project; supervision of data and statistical analyses; acquisition of funding; writing of the manuscript

## A Supplementary Information

**Table S1:** Exon/intron structure of *HMA4* genes and classification of gene parts in the investigated genes from *A. halleri* as well as the control genes from *A. thaliana* and *A. lyrata*.

| Gene part | Length [bp] | Classification |
| --- | --- | --- |
| Exon 0 | 154 (*HMA4-1*), 155 (*HMA4-2*, *HMA4-3* & *A. lyrata*),<br>192 (*A. thaliana*) | 5' UTR |
| Intron I | 327-543 | Intermediate intron |
| Exon I | 213 | Coding exon |
| Intron II | 1123-3035 | Long intron |
| Exon II | 258 | Coding exon |
| Intron III | 1463-3555 | Long intron |
| Exon III | 99 | Coding exon |
| Intron IV | 122-139 | Short intron |
| Exon IV | 255 | Coding exon |
| Intron V | 87-89 | Short intron |
| Exon V | 138 | Coding exon |
| Intron VI | 82-84 | Short intron |
| Exon VI | 330 | Coding exon |
| Intron VII | 83-85 | Short intron |
| Exon VII | 201 | Coding exon |
| Intron VIII | 83-85 | Short intron |
| Exon VIII | 204 | Coding exon |
| Intron IX | 79-85 | Short intron |
| Exon IX | 2127 (*A. thaliana*), 2340 (*A. lyrata*) | Coding exon |
| Exon IXa | 1389, 1401 | Coding exon |
| Intron X | 105-118 | Short intron |
| Exon IXb | 596-612 (378-393 coding) | Coding exon & 3' UTR |

**Table S2:** Pairwise comparisons of diversities and divergences of long introns from Tukey’s HSD test.

| comparison | diff | lwr | upr | p adj |
| --- | --- | --- | --- | --- |
| 1_3-1_2 | -0.015092 | -0.017178 | -0.013005 | 0.000000 |
| 2_3-1_2 | -0.058476 | -0.060509 | -0.056442 | 0.000000 |
| within_HMA4-1-1_2 | -0.052859 | -0.055619 | -0.050099 | 0.000000 |
| within_HMA4-2-1_2 | -0.051188 | -0.053743 | -0.048632 | 0.000000 |
| within_HMA4-3-1_2 | -0.050391 | -0.052946 | -0.047835 | 0.000000 |
| 2_3-1_3 | -0.043384 | -0.045418 | -0.041350 | 0.000000 |
| within_HMA4-1-1_3 | -0.037767 | -0.040528 | -0.035007 | 0.000000 |
| within_HMA4-2-1_3 | -0.036096 | -0.038652 | -0.033541 | 0.000000 |
| within_HMA4-3-1_3 | -0.035299 | -0.037854 | -0.032743 | 0.000000 |
| within_HMA4-1-2_3 | 0.005617 | 0.002896 | 0.008337 | 0.000000 |
| within_HMA4-2-2_3 | 0.007288 | 0.004775 | 0.009801 | 0.000000 |
| within_HMA4-3-2_3 | 0.008085 | 0.005573 | 0.010598 | 0.000000 |
| within_HMA4-2-within_HMA4-1 | 0.001671 | -0.001459 | 0.004801 | 0.645667 |
| within_HMA4-3-within_HMA4-1 | 0.002468 | -0.000661 | 0.005598 | 0.213856 |
| within_HMA4-3-within_HMA4-2 | 0.000797 | -0.002154 | 0.003748 | 0.971838 |

**Table S3:**
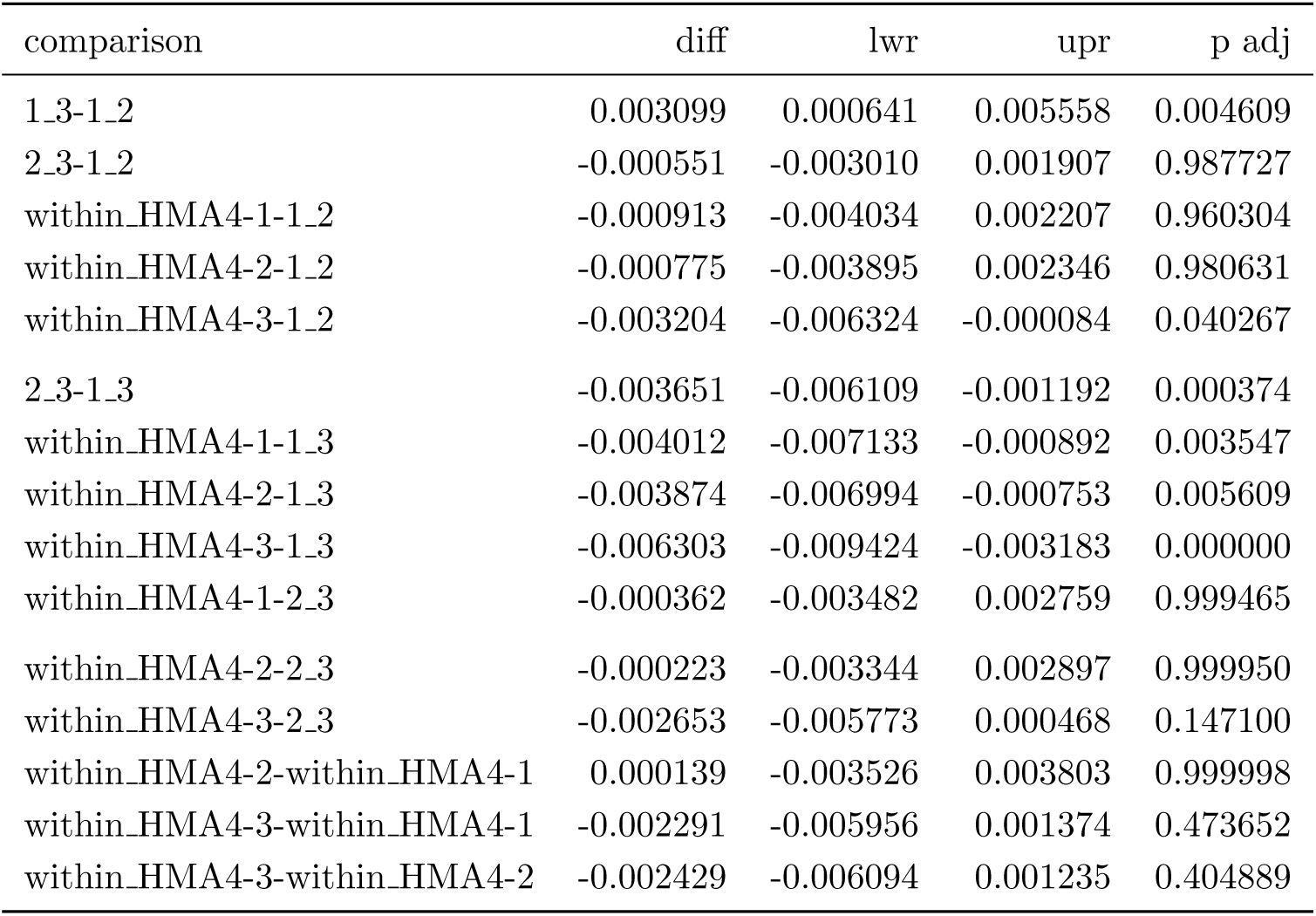
Pairwise comparisons of diversities and divergences of short introns from Tukey’s HSD test.

| comparison | diff | lwr | upr | p adj |
| --- | --- | --- | --- | --- |
| 1_3-1_2 | 0.003099 | 0.000641 | 0.005558 | 0.004609 |
| 2_3-1_2 | -0.000551 | -0.003010 | 0.001907 | 0.987727 |
| within_HMA4-1-1_2 | -0.000913 | -0.004034 | 0.002207 | 0.960304 |
| within_HMA4-2-1_2 | -0.000775 | -0.003895 | 0.002346 | 0.980631 |
| within_HMA4-3-1_2 | -0.003204 | -0.006324 | -0.000084 | 0.040267 |
| 2_3-1_3 | -0.003651 | -0.006109 | -0.001192 | 0.000374 |
| within_HMA4-1-1_3 | -0.004012 | -0.007133 | -0.000892 | 0.003547 |
| within_HMA4-2-1_3 | -0.003874 | -0.006994 | -0.000753 | 0.005609 |
| within_HMA4-3-1_3 | -0.006303 | -0.009424 | -0.003183 | 0.000000 |
| within_HMA4-1-2_3 | -0.000362 | -0.003482 | 0.002759 | 0.999465 |
| within_HMA4-2-2_3 | -0.000223 | -0.003344 | 0.002897 | 0.999950 |
| within_HMA4-3-2_3 | -0.002653 | -0.005773 | 0.000468 | 0.147100 |
| within_HMA4-2-within_HMA4-1 | 0.000139 | -0.003526 | 0.003803 | 0.999998 |
| within_HMA4-3-within_HMA4-1 | -0.002291 | -0.005956 | 0.001374 | 0.473652 |
| within_HMA4-3-within_HMA4-2 | -0.002429 | -0.006094 | 0.001235 | 0.404889 |

**Table S4:** Pairwise comparisons of diversities and divergences of synonymous sites from Tukey’s HSD test.

| comparison | diff | lwr | upr | p adj |
| --- | --- | --- | --- | --- |
| 1_3-1_2 | 0.000380 | -0.000141 | 0.000900 | 0.295895 |
| 2_3-1_2 | -0.002007 | -0.002528 | -0.001486 | 0.000000 |
| within_HMA4-1-1_2 | -0.003588 | -0.004249 | -0.002927 | 0.000000 |
| within_HMA4-2-1_2 | -0.002279 | -0.002940 | -0.001618 | 0.000000 |
| within_HMA4-3-1_2 | -0.003235 | -0.003896 | -0.002574 | 0.000000 |
| 2_3-1_3 | -0.002387 | -0.002907 | -0.001866 | 0.000000 |
| within_HMA4-1-1_3 | -0.003968 | -0.004629 | -0.003307 | 0.000000 |
| within_HMA4-2-1_3 | -0.002658 | -0.003319 | -0.001997 | 0.000000 |
| within_HMA4-3-1_3 | -0.003615 | -0.004276 | -0.002954 | 0.000000 |
| within_HMA4-1-2_3 | -0.001581 | -0.002242 | -0.000920 | 0.000000 |
| within_HMA4-2-2_3 | -0.000272 | -0.000933 | 0.000389 | 0.848097 |
| within_HMA4-3-2_3 | -0.001228 | -0.001889 | -0.000567 | 0.000002 |
| within_HMA4-2-within_HMA4-1 | 0.001310 | 0.000533 | 0.002086 | 0.000028 |
| within_HMA4-3-within_HMA4-1 | 0.000353 | -0.000423 | 0.001129 | 0.783856 |
| within_HMA4-3-within_HMA4-2 | -0.000957 | -0.001733 | -0.000180 | 0.006149 |

**Table S5:** Pairwise comparisons of diversities and divergences of nonsynonymous sites from Tukey’s HSD test.

| comparison | diff | lwr | upr | p adj |
| --- | --- | --- | --- | --- |
| 1_3-1_2 | 0.000394 | 0.000228 | 0.000559 | 0.000000 |
| 2_3-1_2 | 0.000045 | -0.000121 | 0.000210 | 0.972394 |
| within_HMA4-1-1_2 | -0.000595 | -0.000805 | -0.000384 | 0.000000 |
| within_HMA4-2-1_2 | -0.000396 | -0.000606 | -0.000185 | 0.000002 |
| within_HMA4-3-1_2 | -0.000147 | -0.000357 | 0.000064 | 0.347229 |
| 2_3-1_3 | -0.000349 | -0.000515 | -0.000183 | 0.000000 |
| within_HMA4-1-1_3 | -0.000988 | -0.001199 | -0.000778 | 0.000000 |
| within_HMA4-2-1_3 | -0.000789 | -0.000999 | -0.000579 | 0.000000 |
| within_HMA4-3-1_3 | -0.000540 | -0.000751 | -0.000330 | 0.000000 |
| within_HMA4-1-2_3 | -0.000639 | -0.000850 | -0.000429 | 0.000000 |
| within_HMA4-2-2_3 | -0.000440 | -0.000651 | -0.000230 | 0.000000 |
| within_HMA4-3-2_3 | -0.000191 | -0.000402 | 0.000019 | 0.099191 |
| within_HMA4-2-within_HMA4-1 | 0.000199 | -0.000048 | 0.000446 | 0.192718 |
| within_HMA4-3-within_HMA4-1 | 0.000448 | 0.000201 | 0.000695 | 0.000005 |
| within_HMA4-3-within_HMA4-2 | 0.000249 | 0.000002 | 0.000496 | 0.047279 |

**Figure S1:**
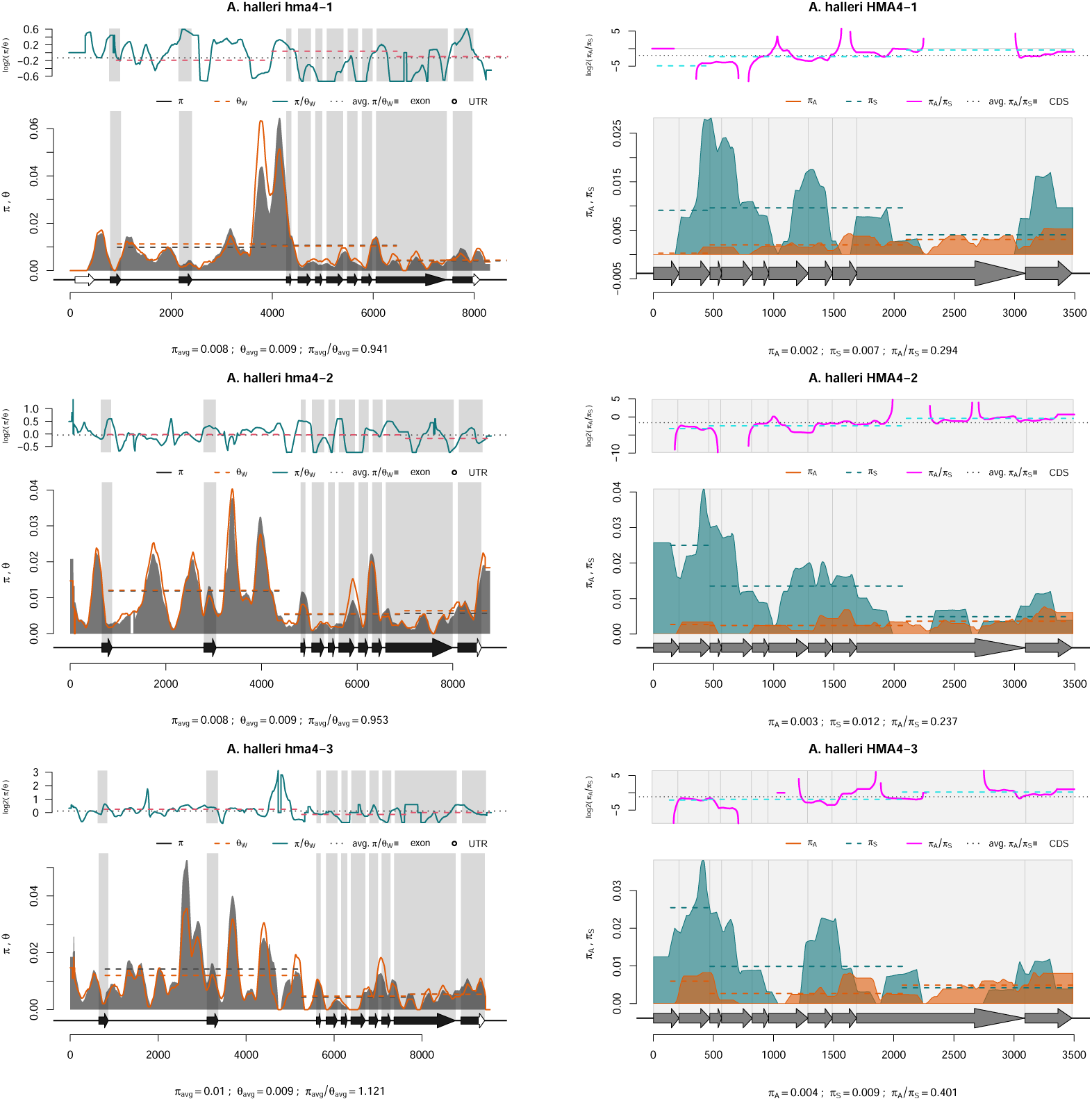
Genetic diversity profiles of *HMA4* genes in *A. halleri*.

**Figure S2:**
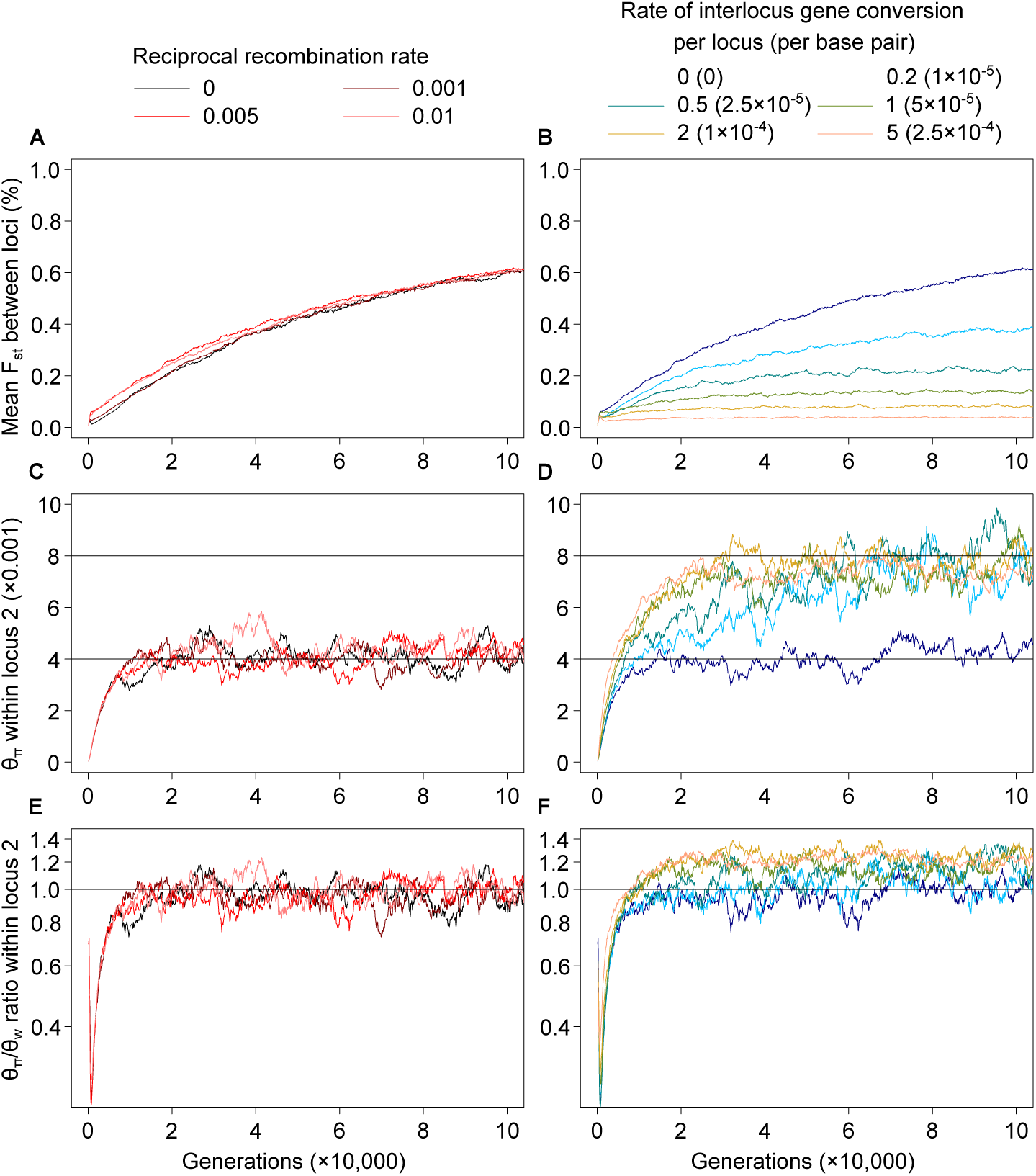
Additional population genetic statistics under a two-locus neutral model without IGC (left side) and with different levels of IGC (right side). Reciprocal recombination rate between loci *r*_rec_ = 0, 0.001, 0.005 or 0.01 for the no-IGC scenarios, and is constant at 0.005 for the IGC scenarios. (A,B) Site-mean *F_st_* values between loci 1 and 2 over 200k generations. (C,D) The value of *θ_π_*, i.e. pairwise nucleotide diversity within locus 2; horizontal line at *θ_π_* = 0.004 per base pair is the single-locus expectation. (E,F) The *θ_π_/θ_w_* ratio for locus 2. This ratio is expected to be 1 (grey line) in an equilibrium population; a ratio greater than 1 is equivalent to a positive Tajima’s *D* statistic and vice versa.

**Figure S3:**
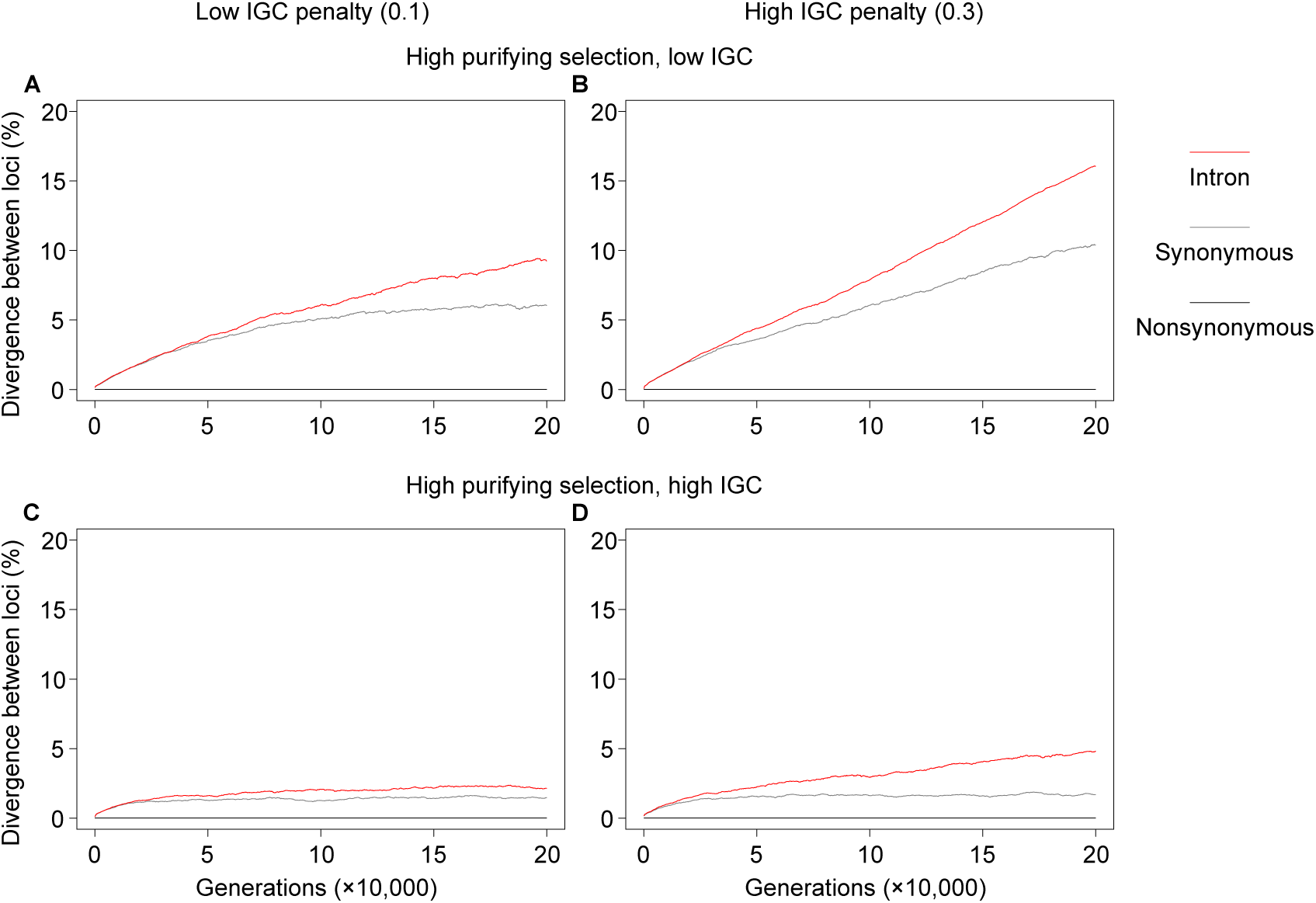
Mean sequence divergence between two loci evolving under selective constraint and a rate of IGC dependent on divergence. The sequence is divided into nonsynonymous positions (black), synonymous positions (grey) and introns (red). Columns: penalty *λ* is low (0.1; A, C) or high (0.3; B, D). Rows: IGC is low (*R*_GC_ = 0.2; A, B) or high (*R*_GC_ = 1.0; C, D). Purifying selection is strong (*s_p_* = 0.01).

## Notes

### Competing Interest Statement

The authors have declared no competing interest.

### Summary of Updates

updated sequence annotation and ensuing sequence analysis; updated Introduction and Discussion; update Fig 4

